# Novel insights into expansion and functional diversification of MIR169 family in tomato

**DOI:** 10.1101/801969

**Authors:** Sombir Rao, Sonia Balyan, Sarita Jha, Saloni Mathur

## Abstract

MIR169 family is an evolutionarily conserved miRNA family in plants. A systematic in-depth analysis of MIR169 family in tomato is lacking. We report eighteen miR169 precursors, annotating new loci for MIR169a, b and d, as well as four novel mature isoforms (MIR169f/g/h/i). The family has expanded by both tandem- and segmental-duplication events during evolution. A tandem-pair ‘MIR169b/b-1 and MIR169b-2/h’ is polycistronic in nature coding for three MIR169b isoforms and a new variant miR169h, that is evidently absent in the wild relatives *S. pennellii* and *S. pimpinellifolium*. Seven novel miR169 targets including RNA-binding-protein, protein-phosphatase, aminotransferase, chaperone, tetratricopeptide-repeat-protein, and transcription factors ARF-9B and SEPELLATA-3 were established by efficient target cleavage in presence of specific precursors as well as increased target abundance upon miR169 chelation by short-tandem-target-mimic construct in transient assays. Comparative antagonistic expression profiles of MIR169:target pairs suggest MIR169 family as ubiquitous regulator of various abiotic stresses (heat, cold, dehydration and salt) and developmental pathways. This regulation is partly brought about by acquisition of new promoters as demonstrated by promoterMIR169:GUS-reporter assays as well as differential processivity of different precursors and miRNA cleavage efficiencies. Thus, the current study augments the functional horizon of MIR169 family with applications for stress tolerance in crops.

**Highlight:** Expansion of MIR169 members by duplication and new mature forms, acquisition of new promoters, differential precursor-miRNA processivity and engaging novel targets increases the functional diversification of MIR169 in tomato. (29/30)

## Introduction

MicroRNAs (miRNAs) are a class of small sequence-specific regulatory RNAs that mediate mRNA cleavage, translational repression and methylation of their target genes. They have essential roles in plant growth, development and stress tolerance (Reinhart et al. 2002; Bartel et al. 2004; Guan et al. 2013; Sorin et al. 2014; Hivrale et al. 2016; Bai et al. 2018). While sequencing technology advancement has heralded an era of identification of a large number of plant miRNAs, only a few of them have been characterized functionally. For instance, 25 conserved miRNA families and 15 novel miRNAs responding to pathogen have been identified in maize (Thiebaut et al. 2014), nitrogen deficiency-responsive miRNAs were discovered in both Arabidopsis (Liang et al. 2012) and maize (Zhao et al. 2012), phosphate deficiency-responsive miRNAs were studied in soybean (Xu et al. 2013), heat-responsive miRNAs were analysed in *Betula luminifera* (Ying et al. 2017), banana (Zhu et al. 2019) and wheat (Ravichandran et al. 2019), drought-responsive unique miRNAs were defined in rice (Kansal et al. 2015; Mutum et al. 2016), drought and rust infection stress-responsive miRNAs were reported in soybean (Kulcheski et al. 2011) among others.

Many plant miRNAs are evolutionarily conserved among species and can be grouped into distinct families having one or more genomic loci. Within a miRNA family, even though precursors of different members produce the same or similar mature miRNAs, they exhibit differential spatial and temporal expression profiles creating functional diversification, leading to the modulation of distinct target genes (Baker et al. 2005). MIR169 has been identified in more than forty species (Sunkar & Jagadeeswaran, 2008), where it often makes up the largest miRNA family. The Arabidopsis miR169 family contains 14 genes, rice has 18 members and Sorghum MIR169 family is 17 members large (http://www.mirbase.org/). However, all precursors do not mature into as many miRNAs, for example, in Arabidopsis only four mature miR169 isoforms (a, bc, defg and hijklmn), differing by one or two nucleotides at the 5′ or 3′ ends are produced from the 14 precursors (Li et al. 2010). Ubiquitous presence of MIR169 family across genera suggests its important regulatory roles.

The Nuclear Factor-YA (NF-YA) family members, are established as the main targets of miR169 family and regulate plant response towards abiotic and biotic stress mainly via ABA pathway (Ding, Zeng & He, 2016; Luan et al. 2015; Zhao et al. 2009). Overexpression of miR169c in tomato provides dehydration tolerance by regulating stomatal aperture through post transcriptionally regulating Multi-Drug Resistant protein-1 (Zhang et al. 2010, 2011). Arabidopsis seedlings show cold stress induced over-accumulation of miR169 that correlates with a reduction of some NF-YA target transcripts (Zhou et al. 2008; Lee et al. 2010). miR169 may also play a role in long-distance signaling, as its abundance in phloem sap decreases during nitrogen (N) and phosphorus (P) limitation (Pant et al. 2009; Buhtz et al. 2010) and several miR169 NF-YA targets are induced (Pant et al. 2009; Zhao et al. 2011). Overexpression of miR169 in Arabidopsis mediates early flowering in response to stress and regulates root architecture by repressing the AtNF-YA2 transcription factor (Xu et al. 2014; Sorin et al. 2014). In *Medicago truncatula* miR169-mediated restriction of MtHAP2-1 expression to the nodule meristematic zone is essential for the differentiation of nodule cells (Combier et al. 2006). In addition, miR169a and miR169i have been reported to regulate tolerance to drought stress through targeting NF-YA5 mRNA in Arabidopsis (Li et al. 2008; Du et al. 2017). Osa-miR169g and osa-miR169n/o was upregulated by high salinity, and selectively cleave NF-YA8 in response to high salinity stress (Zhao et al. 2009). Overexpression of miR169a results in hyper susceptibility to different *Magnaporthe oryzae* strains (Li et al. 2017). Overexpression of miR169o leads to susceptibility to bacterial blight in rice (Yu et al. 2018) while in tomato miR169 enhances resistance to gray mold (Li et al. 2016; Gu et al. 2010).

To understand functional diversification of this important miRNA family in tomato, we have done a comprehensive analysis to identify MIR169 family members. Further, expression analysis of MIR169 family members in different abiotic stresses and developmental stages revealed its tissue-specific and stress-responsive nature in a redundant as well as specific manner. We highlight the yet unexplored roles of miR169 family in different heat stress regimes. MIR169 promoter:GUS-reporter assays suggest acquisition of new promoters during evolution that provides functional diversification. This study identifies and validates novel targets of the family in tomato and explores their roles in different abiotic stresses and tissues.

## Materials and Methods

### Plant material and stress conditions

Three-days-old seedlings of tomato cultivar, CLN1621L (CLN) of uniform growth were transplanted in plastic pots, filled with soilrite and placed in a plant growth chamber (CMP6050, Conviron, Canada) maintained at 26 °C/21 °C (day/night: 16/8 h), relative humidity 60%, light intensity 300 µM per m^2^ per sec. For collection of flower and fruit tissues plants were grown up to 80-90 days while thirty-days-old plants were subjected to all stresses (different heat stress regimes, cold, dehydration and salt). Heat stress consisted of the following regimes: Basal stress: Plants were directly exposed for 4.5 h at 45 °C. Acquired heat stress: Plants were initially exposed for 4 h at a gradual increasing range of temperature from 26 °C to 45 °C then exposed additionally for 4.5 h at 45 °C. In addition, plants from the second regime were allowed to recover overnight in a chamber maintained at 26 °C/21 °C (day/night: 16/8 h). Leaves from these experimental plants were harvested at relevant time points, frozen in liquid nitrogen and kept at − 80 °C until use. For, low temperature stress, plants were placed at 4±2 °C for 2 days. Dehydration stress was imposed by withholding water for 12 days. For salt stress a set of plants were transferred to half strength MS medium and stabilized for 2 days in a growth chamber, plants were then transferred to nutrient solution containing 170 mM sodium chloride and the nutrient solution without sodium chloride, served as control. The concentrations of solutions used were chosen based on preliminary experiments (Hu et al, 2009; Sarkar et al, 2009).

### MIR169 gene prediction in tomato

Previously known 799 miRNA169 sequences of diverse plant species were downloaded from miRNA registry database i.e. miRBase (http://www.mirbase.org, Released 22.1: October, 2018). Non-redundant miRNAs were used as reference for finding the homologs in tomato. Homology search of reference miRNAs against the tomato genome (Tomato genome chromosome build SL4.0) was performed at sol genomics network (SGN) database by blastn for short query sequences, by increasing the e-value parameter (E = 10). Hits with ≥ 75% similarity were short-listed. The pre-miRNA sequences were extracted from the region 80-250 nucleotide upstream of the beginning of the mature miRNA to 80-250 nt downstream of the miRNA (Singh and Nagaraju, 2008). The fold-back secondary structures of pre-miRNAs were predicted using RNA fold and M fold (Zuker, 2003). The following criteria were used for selecting the pre-miRNA structures according to Lu and Yang (2010): (1) the sequence should fold into an appropriate stem-loop hairpin secondary structure (2) predicted mature miRNAs with no more than 3 nt substitutions as compared with the known miRNAs (3) no more than 6 mismatches are between the predicted mature miRNA sequence and its opposite miRNA* sequence in the secondary structure (4) mature miRNA should be in the stem region of the hairpin structure (5) no loop or break is in the miRNA* sequences, and (6) predicted secondary structure has high Minimum Folding Energy Index (MFEI > 0.85) and negative Minimum Folding Energy (MFE).

### Experimental validation of predicted MIR169 genes

Two µg of high quality total RNA was used to synthesize cDNA using high capacity cDNA reverse transcription kit (Applied Biosystems, USA) as per the instructions of the manufacturer and described in Paul et al. (2016). All predicted MIR169 precursors were amplified and cloned into TA-vector using InsTAclone PCR Cloning Kit, (Thermo-Fisher Scientific, USA). Cloned precursors were further confirmed by sequencing.

#### Chromosomal localization, duplication and phylogenetic analysis of MIR169 family

To determine the location of MIR169 genes on tomato chromosomes, a BLASTN search with SGN database was conducted. The resulting position of the MIR169 genes on the tomato chromosome was mapped by using Mapchart 2.2 (Voorrips 2002). For phylogeny MIR169 precursor sequences from rice and Arabidopsis were extracted from miRBASE. Multiple alignments of all MIR69 precursor sequences and phylograms were drawn with the ClustalX program using the neighbor joining (NJ) method with 1,000 bootstrap replications. Since non-coding transcribed regions (precursors) were being compared having a small ∼21 bp conserved miRNA within approximately 200 bp of precursor, a bootstrap value of at least 40% was used to delineate clades. Phylogenetic tree was visualized using Figtreev1.

### Expression analysis of MIR169 family members and targets

Total RNA was isolated from leaf tissues using Ribozol (Ameresco, USA), whereas from flower and fruit, RNA was isolated as described by Muoki et al. (2012b). DNase I (Ambion, USA) treated total RNA was used to synthesize cDNA using high capacity cDNA reverse transcription kit (Applied Biosystem, USA) following the manufacturer’s protocol. Gene expression was performed on 7900 HT fast real time PCR system (Applied Biosystem, USA) using 2× Brilliant III SYBR^@^ Green qPCR Master Mix (Agilent Technologies, USA) and gene specific primers (Supplementary Table 1). All qPCRs were run with biological and technical replicates. A no-template control was kept to check for contamination under default settings. A final melt-curve analysis was performed (55° to 95 °C) to verify the specificity of amplicons. The raw threshold cycle (Ct) values were normalized against the housekeeping gene *Actin.* The microarray data-based expression profiles of miRNA targets in different organs and abiotic stresses including heat, dehydration, and salt stress condition were assessed using plant biology tool at Genevestigator (http://www.genevestigator.ethz.ch) on Le_10K microarray platform. The expression was computed in form of heat maps for different organs and abiotic stress conditions.

### Prediction of miR169 targets by degradome analysis

The target prediction of miR169 members was done by analyzing the PARE datasets (GSM1047560, GSM1047562, GSM1047563, GSM553688, GSM553689, GSM553690, GSM1379695, GSM1379694, GSM1372427 and GSM1372426) of different tissues of tomato from NCBI (https://www.ncbi.nlm.nih.gov/sra). Briefly, all the PARE datasets was trimmed using Trim adapter tool of CLC Genomics workbench version 9. All the miR169 mature forms were used as miRNA query against the tomato gene model sequences of tomato (ITAG version 4) following the CleaveLand4 pipeline (Addo-Quaye et al. 2009) with the default settings. Further the miRNA/mRNA pairs with an alignment score ≤ 7, total PARE reads more ≥2 being cut at 10^th^ position (+/- 1 base) with evidence from at least 2 datasets were considered as targets.

### Constructs generation

For promoter constructs Pro-SlyNFYA10:GUS, Pro-Sly-MIR169a:GUS, Pro-Sly-MIR169b:GUS, Pro-Sly-MIR169a-1:GUS, Pro-Sly-MIR169a-2:GUS, Pro-Sly-MIR169b:GUS, Pro-Sly-MIR169b-1:GUS, Pro-Sly-MIR169b-3:GUS, Pro-Sly-MIR169b-4:GUS, Pro-Sly-MIR169c:GUS, Pro-Sly-MIR169d:GUS and Pro-Sly-MIR169d-1:GUS, 2000 bp region upstream of the start codon (for target) or precursor start site was amplified from genomic DNA and cloned into pbi101 vector (For primer details refer Supplementary Table 1). For the overexpression of precursors constitutive CaMV 35S promoter was used and named 169a-OX, miR169a-1-OX, miR169b-OX, miR169b-1-OX, miR169c-OX, miR169d-OX, miR169d-1-OX and miR169i-OX, 300–400 bp precursor fragments were amplified from genomic DNA, cloned into pbi121. For the transient experiments for MIR169 target validation, the 3′ UTRs of NF-YA1, NF-YA3, NF-YA8, NF-YA9 and NF-YA10 and CDS of novel targets were fused to the GFP coding region and cloned into the pCAMBIA1302 vector under CaMV 35S promoter.

### Agro-infiltration and transient assay for promoter analysis and miRNA target confirmation

Agro-infiltration was performed in 5-days-old tomato seedlings for GUS aided promoter analysis and 4-week-old *Nicotiana benthamiana* leaves for target validation. Various Pro-Sly-MIR169s: GUS, miR169s-OX and GFP-NF-YAs overexpression constructs were transformed into *Agrobacterium tumefaciens* EHA105 strain. For infiltrations, overnight cultures of individual constructs were harvested and suspended at OD 1 in infiltration buffer (10 mM MgCl2, 10 mM MES, pH 5.8, 0.5% glucose and 150μM acetosyringone) and incubated for 3 h at room temperature. Various reporter cultures were infiltrated in tomato seedlings by vacuum. For target validation, a set of *N. benthamiana* leaves were infiltrated by syringe with the target constructs alone or in a 1:1 ratio of precursor-overexpression:target constructs The seedlings/plants were kept in growth chamber maintained at 26□ C and light intensity of 300 µM per m^2^ per sec and harvested after 2 days for GUS staining and/or RNA extraction For target validation, HPT gene in vector was used to normalize target abundance in qPCR experiments. For precursor efficiency qPCR assays, the level of precursor expression was also normalized by NPTII gene in the OX construct.

### Histochemical GUS Assays

For histochemical GUS localization, samples were incubated for 12 h at 37°C with the substrate solution (1 mM 5-bromo-4-chloro-3-indolyl-β-D-glucuronide, pH 7.0, 100 mM sodium phosphate buffer, 10 mM Na2EDTA, 0.5 mM potassium ferricyanide, 0.5 mM potassium ferrocyanide, and 0.1% Triton X-100). Stained seedlings/leaves were washed with de-staining solution containing ethanol:acetone:glycerol (3:1:1) to eliminate chlorophyll, and photographed with Nikon SMZ1000 Stereomicroscope (Tokyo, Japan). GUS transcript was quantified using qPCR. The assay was performed with three biological replicates.

### Short-tandem-target-mimic (STTM) to sponge miRNA169 in tomato

For STTM169 suppression vector construction, first a cassette containing 2×35S promoter and *NOS* terminator was cloned in pCAMBIA2300 multi-cloning site. Two overlapping long oligos were synthesized as per Tang et al. (2012) paper to make STTM fragment. The oligos were annealed, amplified and inserted downstream of the 2×35S promoter in the modified pCAMBIA2300 vector. The construct was transformed into *Agrobacterium tumefaciens* strain EHA105, and vacuum infiltrated in tomato seedlings. MIRNA169 chelation was confirmed by measuring validated targets by target-specific qRT-PCR.

## Results and Discussion

### An amplified repertoire of MIR169 members in tomato

Presence of miR169 family in diverse plant species including monocots, dicots, gymnosperms and ferns (Michael et al. 2005), highlights the critical roles this miRNA family has in regulating plant development and stress response. Multiple MIR169 family members have been identified in Arabidopsis, sorghum, maize, banana and carrot (Calvino et al. 2013; Mica et al. 2006; Song et al. 2018; Bhan et al. 2019). The current miRBase version22 reports fourteen MIR169 loci in Arabidopsis (a to n) and only five loci for tomato (a to e). This is very striking as tomato genome size is nearly five times that of Arabidopsis genome and the Solanum lineage is reported to have undergone two consecutive genome triplication, one being ancient and one recent (The tomato genome consortium, 2012). One would therefore, expect many more MIR169 loci in tomato. This prompted us to perform a genome-wide survey to identify new MIR169 members in tomato. Using an in-house designed pipeline (Supplementary Figure 1a) we identified 21 putative miRNA169 genomic loci in tomato (Table 1). The length of all predicted MIR169 precursors range from 100 to 192 nucleotides (Table 1), which is in the acceptable range of plant miRNA precursors that are reported from 55 to 930 nucleotides long (Thakur et al. 2011). Furthermore, these precursors also satisfy the criteria used to distinguish miRNAs from all coding and other non-coding RNAs (Zhang et al. 2006).

**Table 1:** Characteristics of the putative pre-MIR169s in tomato. Details of different putative MIR169 loci identified in the study. The start end positions of the predicted precursors on different chromosomes is shown as per SGN build4.0. Chr: Chromosome; Str: Strand; LP: length of pre-miRNA; MFE: minimum fold energy; NM: number of mismatches between predicted miRNA and miRNA*.

| S. No | MIR169 Ids | Chr | Start | end | Str | LP<sup>a</sup> | MFE kcal/mol | NM<sup>b</sup> | GC % |
| --- | --- | --- | --- | --- | --- | --- | --- | --- | --- |
| 1 | Sly-MIR169a | Ch07 | 4195624 | 4195772 | + | 149 | -68.80 | 1 | 42.9 |
| 2 | Sly-miR169a-1 | Ch02 | 48346026 | 48346174 | + | 149 | -61.7 | 0 | 42.9 |
| 3 | Sly-miR169a-2 | Ch07 | 4219875 | 4220022 | + | 148 | -69.60 | 1 | 43.2 |
| 4 | Sly-MIR169b | Ch07 | 2147189 | 2147322 | + | 134 | -49.47 | 2 | 36.5 |
| 5 | Sly-miR169b-1 | Ch07 | 2146991 | 2147094 | + | 104 | -46.33 | 2 | 39.4 |
| 6 | Sly-miR169b-2 | Ch07 | 2153197 | 2153329 | + | 133 | -48 | 2 | 36.8 |
| 7 | Sly-miR169b-3 | Ch03 | 5167831 | 5167977 | + | 143 | -49.10 | 2 | 40.1 |
| 8 | Sly-miR169b-4 | Ch03 | 5174789 | 5174637 | - | 153 | -51.00 | 3 | 39.2 |
| 9 | Sly-MIR169c | Ch07 | 4181057 | 4181203 | + | 147 | -80.5 | 1 | 41.4 |
| 10 | Sly-MIR169d | Ch07 | 2144070 | 2144245 | - | 176 | -72.4 | 1 | 32.3 |
| 11 | Sly-miR169d-1 | Ch08 | 55689928 | 55690067 | + | 140 | -40.79 | 2 | 39.2 |
| 12 | Sly-miR169d-2 | Ch08 | 53593952 | 53594051 | + | 100 | -44.62 | 2 | 45 |
| 13 | Sly-miR169d-3 | Ch08 | 55698703 | 55698818 | + | 116 | -40.03 | 1 | 31.8 |
| 14 | Sly-miR169d-4 | Ch08 | 55797862 | 55797981 | + | 121 | -43.15 | 1 | 32 |
| 15 | Sly-miR169d-5 | Ch03 | 54665334 | 54665471 | + | 138 | -41.84 | 3 | 30 |
| 16 | Sly-MIR169e | Ch08 | 55684858 | 55685040 | + | 192 | -78.4 | 3 | 38 |
| 17 | Sly-MIR169f | Ch01 | 2525744 | 2525913 | + | 170 | -41.53 | 5 | 35.8 |
| 18 | Sly-MIR169g | Ch06 | 42401751 | 42401894 | + | 144 | -42.8 | 4 | 34.7 |
| 19 | Sly-MIR169h | Ch07 | 2153624 | 2153753 | + | 130 | -43.2 | 1 | 43.8 |
| 20 | Sly-MIR169i | Ch07 | 4104944 | 41105067 | + | 124 | -61.1 | 1 | 43.5 |
| 21 | Sly-MIR169j | Ch01 | 42129523 | 42129661 | + | 138 | -52.78 | 6 | 28 |

We have validated eighteen of these members by cloning and sequencing of amplicons derived from a pool of cDNA representing different plant tissues as well as stress samples (Supplementary Figure 1b). These miR169 loci include the five known and thirteen new members. We report two new precursors for Sly-miR169a, four for Sly-miR169b and 3 new precursors for Sly-miR169d. Additionally, 4 new precursors were validated which produce four different sequence variants of mature Sly-miR169 (f to i). This increases the mature Sly-miR169 count from five to nine (Supplementary Figure 1b-c).

Seven of these mature forms are 21mers while Sly-miR169e and Sly-miR169f are 22- and 20 nucleotide long, respectively. The mature Sly-miR169s exhibit similarity in size and sequence with other known plant paralogs (Supplementary Figure1c). The nine mature Sly-MIR169 forms are 75% identical over their length having 7 identical bases in all (Supplementary Figure 1c). Six mature sequence variants of miR169 have ‘T’ as 5’ terminal nucleotide suggesting their loading primarily on AGO1, these include three of the novel forms (f, g and h). In contrast, three miR169 mature variants have ‘C’ as 5’ terminal base (including one novel form, i) suggesting their loading majorly on AGO5 followed by AGO1 (Mi et al. 2008). This differential AGO-mediated loading of different miR169 isoforms could provide functional diversity to different miRNA variants under different spatio-temporal contexts or stress responses.

### Evolutionary diversification of MIR169s in tomato

MicroRNAs tend to evolve continuously by accumulating sequence variations to recognize newer distinct targets and attain functional divergence. It is noteworthy, that in comparison to 21 MIR169 loci in cultivated tomato *S. lycopersicum* (Sly), there are 16 loci in the closest wild relative *S. pimpinellifolium* (Spi) and 20 members in distant wild relative *S. pennellii* (Spe) (Supplementarry Table 2). We find that out of the four novel Sly mature forms identified in our study, forms ‘f and g’ are absent in Spe but are present in Spi. Further, the mature form ‘h’ is absent in both wild relatives while ‘i and j’ forms are present in Spe and the cultivated variety. Thus, among the analysed tomato lineage, the observed variation in copy number of a miRNA169 form between cultivated and wild varieties could have a role in miRNA dosage effect, as is reported for other miRNAs (Marcinkowska et al. 2011). Moreover, new mature variants that are present in the cultivated tomato expand the possibility of engaging new targets which may have roles in tomato domestication.

Further, to understand the evolutionary divergence of MIR169 family in tomato, we performed a phylogenetic analysis with two model plants, the dicot Arabidopsis and monocot rice. The phylogram of fifty MIR169 precursors from tomato, Arabidopsis and rice forms four major clades (Figure 1a). Nearly 86% (43/50) of MIR169 members from the three plant species group into two clades (Clade II and IV) signifying a common ancestry and similar course of evolutionary path. However, dicot-specific clade I and sub-clade IV-A as well as monocot-specific clade III and sub-clade IV-B point to further lineage-specific diversification as these plants evolved to gain functional distinction. Interestingly, Ath-MIR169a does not form part of any specific clade and appears to have diverged independently. Further, multiple sequence alignment of mature miR169s from tomato, rice and Arabidopsis strongly supports the conserved property of miR169 family (Figure 1b). Only, Sly-miR169f form clusters with two rice MIR169s viz., miR169f.2 and miR169i in a separate group and shows less sequence conservation with other forms. The majority of miR169 sequences (49 out of 52) are highly conserved between position 2-21 and diversity lies in their ends. The 5’ variability may have bearing on association with different AGOs (Mi et al. 2008) and the 3′ end may act as a key determinant regulating miRNA activity and stability (Sheu-Gruttadauria et al. 2019).

**Figure 1:**
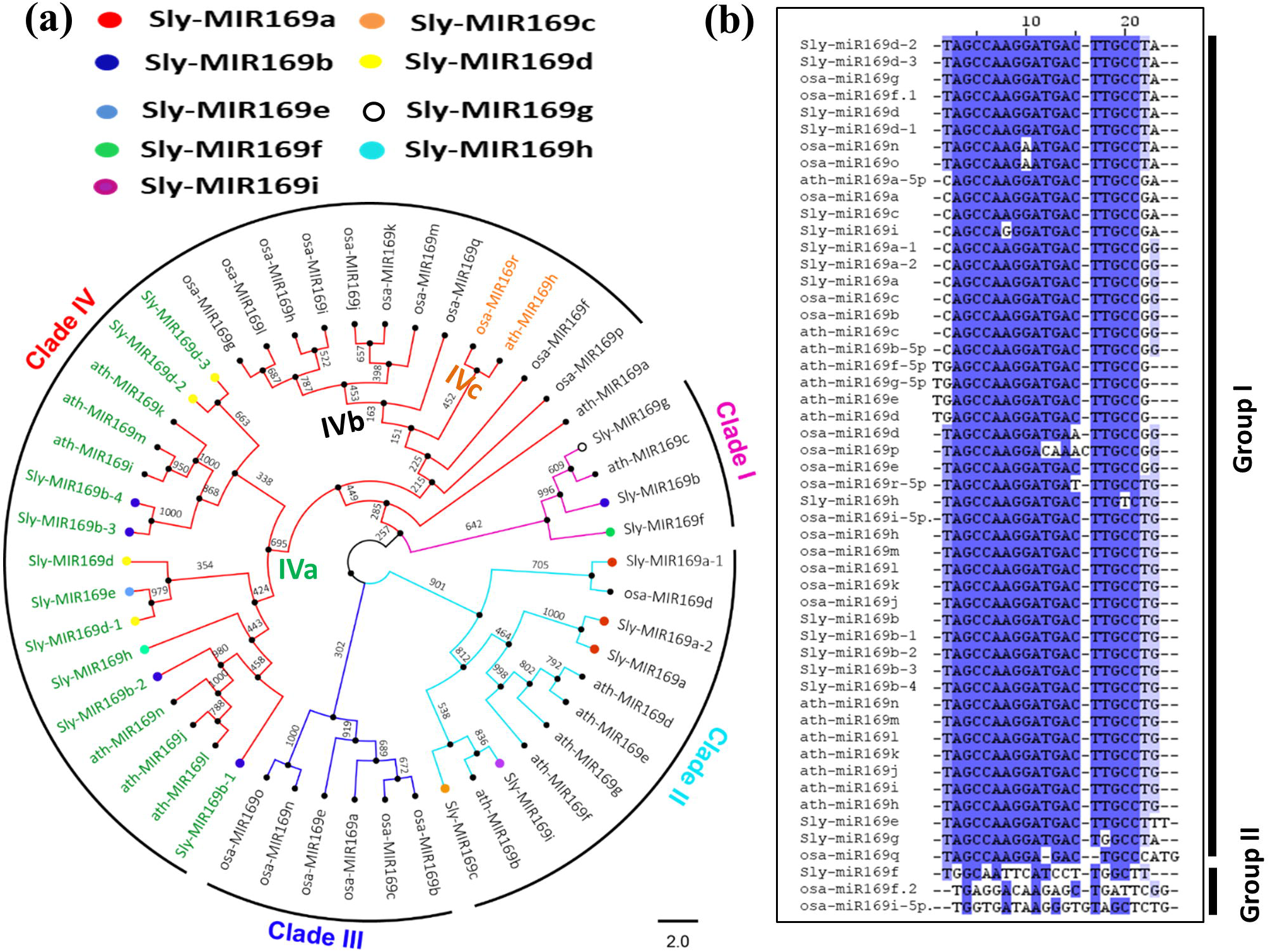
Phylogenetic analysis and alignment of miR169 family members in tomato, Arabidopsis and rice. (a) An unrooted tree of miR169 precursor sequences from tomato, Arabidopsis and rice using Neighbour-Joining method. The bootstrap support from 1,000 replicates are shown at the base of nodes. The major clades are marked by arcs and numbered. Clade IV is further divided into three sub-clades marked at the base of node by different colored text. Tomato-specific MIR169 members are highlighted by different colored dots; precursors coding for same mature form have been given identical colors. The scale bar represents the nucleotide substitution per site. (b) Multiple sequence alignment of mature miR169s of tomato, Arabidopsis and rice, to show the degree of conservation among family members. Identical bases in all sequences are marked in dark blue.

### Expansion of MIR169 family by duplication events in tomato

Segmental and tandem duplication events as well as transposition are considered as the main sources for gene family expansion within plant genomes (Cannon et al. 2004). Localization of the 18 tomato MIR169 genes shows four gene clusters (Supplementary Figure S4) that are scattered on chromosome 3, −7 and −8. It is reported that nearly 20% of plant miRNAs are clustered, and generally contain conserved miRNA members of the same family (Merchan et al. 2009). Gene cluster MIR169b-3/b-4 is a tandem duplication that is located within 6 kb DNA segment on chromosome 3 (Table1, Supplementary Figure S4), has 95% sequence similarity and share similar precursor stem-loop structures (Supplementary Figures S2a, S3). This paralogous pair clusters at 1000 bootstrap iteration in a tomato-specific MIR169 phylogram (Supplementary Figure S2b). Two other tandemly duplicated pairs viz., Sly-MIR169b/h and Sly-miR169b-1/b2 are positioned 6.5 kb and 7 kb apart, respectively, on chromosome 7. The cluster Sly-MIR169a/a-2 groups together at 1000 bootstrap value, exhibits 89% sequence similarity and similar stem-loop structures (Supplementary Figures S2a, b, S3). This pair has evolved by a segmental duplication event, the genes being ∼24kb apart on chromosome 7. Expansion by duplication for miR156, miR159, miR166, and miR395 has been reported in Arabidopsis (Maher et al. 2006), for miR395 in rice (Guddeti et al. 2005) and for miR166 in soybean (Li et al. 2017). In tomato, both tandem and segmental duplications appear to have expanded the MIR169 family during the course of evolution.

### Tomato MIR169b-1/b and MIR169b-2/h are polycistronic and the latter is specie-specific

Genome wide studies on plant miRNA genes have shown that many clustered miRNAs can be transcribed simultaneously into a single polycistronic unit (Bartel 2004, Tanzer et al. 2004, Altuvia et al. 2005, Salvador-Guirao et al. 2018). Among the four tomato MIR169 clusters, the precursors of MIR169b-1 and MIR169b are 92bp apart and the distance between MIR169b-2 and MIR169h precursors is 297bp. This close proximity of their precursors indicates that they may be transcribed together as a single unit (Supplementary Figure S5a). To confirm this, primer pairs spanning the stem-loop regions of adjacent miRNA precursor pairs in question were used to detect amplicons from primary miRNA transcripts. Amplification of specific 280bp and 380bp fragments followed by sequence confirmation for MIR169b-1/b and MIR169b-2/h, respectively, corroborates their polycistronic nature (Supplementary Figure S5c). It is noteworthy, that out of the four miR169s, three code for the ‘b’ form and one is the new variant ‘h’ that is present only in the cultivated tomato (Supplementary Figure S6b, Supplementary Table 2). We therefore, extended the study to find whether these duplications (MIR169b-1/b and MIR169b-2/h) are conserved between cultivated and wild tomato lineages. Comparative sequence analysis of both pair shows that the two precursors are in close proximity in both wild relatives also, suggesting an ancient conserved polycistronic feature (Supplementary Figure S7, S8). However, structural rearrangements during evolution have resulted in variable inter-precursor length, for MIR169b-1/b it is 94 bp in both Spi and Sly while a 9 bp deletion was observed in Spe making the inter-precursor length 85 (Supplementary Figure S7). Furthermore, MIR169b-2/h pair inter-precursor length is 279 bp, 337 bp and 294 bp in Spe, Spi and Sly, respectively (Supplementary Figure S6a, S8). Several sequence and structural determinants have been reported to play significant roles in precursor processing and mature miRNA production (Chorostecki et al. 2017; Jagtap et al. 2014). It would be tempting to study the relevance of this variability in length as well as sequence, on precursor processing by the Dicer-like protein and other associated protein complex.

### Regulatory diversity of duplicated MIR169 genes by different promoter acquisition

We then asked the question whether duplication of MIR169s in tomato has created any regulatory diversity as has been reported for other plant miRNAs (Maher et al. 2006; Sieber et al. 2007). We hooked the promoters of the duplicated Sly-MIR169 pairs to GUS reporter gene and visualized its expression in tomato seedlings. We find that while pro-Sly-MIR169a:GUS expresses strongly in the whole seedling, its paralog pro-Sly-MIR169a-2:GUS does not (Supplementary Figure S9a). We have shown that Sly-MIR169b1/b2 and Sly-MIR169b/h are tandem duplicates but are transcribed as Sly-MIR169b1/b and Sly-MIR169b2/h polycistronic units (previous sections). The promoter of Sly-MIR169b-2/h shows strong transcriptional activity in seedlings but that of Sly-MIR169b-1/b does not (Supplementary Figure S9c). Both the promoters of the third paralogous pair Sly-MIR169b-3 and Sly-MIR169b-4 on the other hand does not show any GUS expression in tomato seedlings (Supplementary Figure S9b). Thus, duplication has allowed newly formed loci to acquire different promoters contributing to specific spatial expression that could be pivotal for regulating target levels. Further, comparative study of promoter of these duplicated pairs shows differences in both numbers as well as types of cis-elements (Supplementary table 3) that may impart regulatory diversity to different MIR169s in stress responsiveness and/or plant development.

### MIR169 has roles in abiotic stresses

The miR169 family is known to play significant roles in stress responses like drought, salt, ABA and heat in plants (Zhao et al. 2007; Zhao et al. 2009; Li et al. 2010). We evaluated the expression pattern of all the tomato MIR169 genes in response heat stress (HS), salinity, dehydration and low temperature (Figure 2a) by qRT-PCR. All Sly-MIR169 genes except Sly-MIR169a and Sly-MIR169d-1 express at very low levels (exhibiting high Ct-values) in control conditions in tomato leaves (Supplementary table 4). However, all sixteen tomato MIR169 genes exhibit significant differential (up-/down-) expression in response to one or the other abiotic stresses (Figure 2a, b). There is up-regulation of 11, 15, 3 and 7 precursors and down-regulation of 1, 0, 12 and 3 precursors in low temperature, HS, dehydration, and salinity stress, respectively (Figure 2b). The promoters of these stress-responsive MIR169s harbor dehydration responsive elements, low temperature responsive cis-elements and heat stress elements (Supplementary Table S3). Nine MIR169s are uniquely down-regulated in dehydration response while one MIR169 is uniquely down-regulated in cold stress only (Figure 2b). Only Sly-MIR169f is not up-regulated in response to any stress. Three MIR169s (Sly-MIR169b-1/b, Sly-MIR169b-4 and Sly-MIR169c) are up-regulated in all abiotic stresses and, four MIR169s are commonly up-regulated in cold, heat and salt stresses (Figure 2b). Our findings show that different MIR169 precursors are stress responsive; over-expression of these miRNAs has potential for engineering multiple stress tolerance in crop plants.

**Figure 2:**
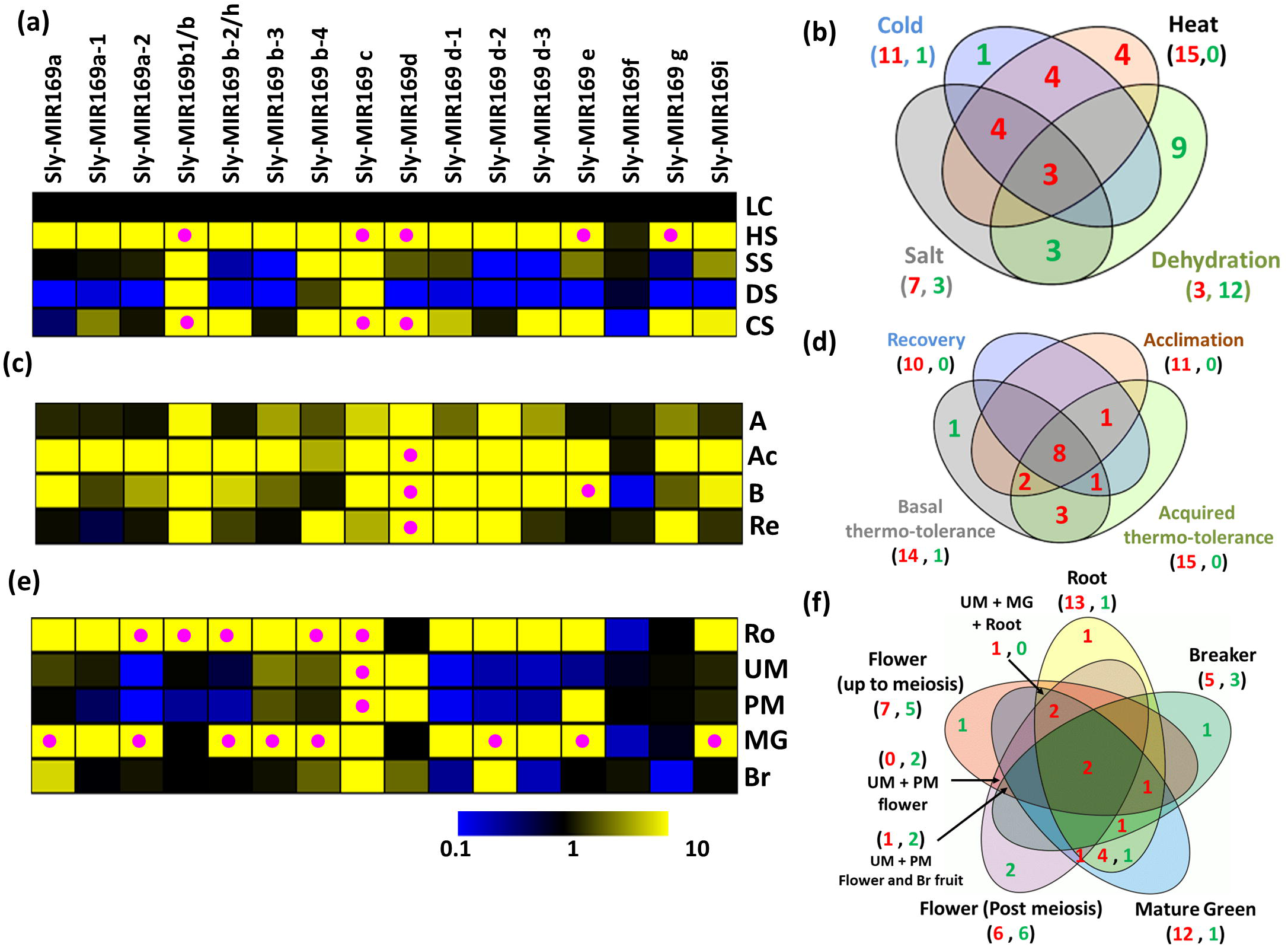
Expression profiles of tomato MIR169s precursors in abiotic stress and different plant organs by qRT-PCR: (a) Heat map of expression of MIR169 members in response to different abiotic stresses. LC: Leaf control, HS: Heat stress, SS: Salt stress, DS: Dehydration stress, CS: Cold stress. (b) Venn diagrams of up-(red) and down-regulated (green) genes in different abiotic stresses. (c) Heat map of MIR169s expression profiles in leaf in response to different heat stress regimes. A: Acclimation, Ac: Acquired thermotolerance, B: Basal thermotolerance, Re: recovery. (d) Venn diagrams of up-(red) and down-regulated (green) genes in different heat stress regimes and heat stress recovery. (e) Expression patterns of MIR169s in different plant organs normalized with control leaf. Ro: root, UM: Up-to meiosis bud pool, PM: post meiosis bud pool, MG: mature green fruit, Br: breaker fruit. (f) Venn diagrams of up-(red) and down-regulated (green) genes in different organs of tomato plant. Expression levels of actin was used as internal references for expression of Sly-MIR169s. Values presented in the heat maps were means of independent experiments. Genes expressing more than 50-folds are depicted by pink dots in (a), (c) and (e).

Since fifteen out of the sixteen MIR169 loci were highly expressed in HS, we further explored the variation of MIR169 response to different HS regimes as well as HS recovery. The regimes include basal HS (directly exposed to 45°C), acclimation stress (gradual increase in temperature) and acquired HS (an acclimation phase prior to harsh HS at 45°C). It is noteworthy, that fourteen of the fifteen HS-responsive precursors are strongly up-regulated (≥10 fold) in at-least one of the HS regimes (Figure 2c, Supplementary Table S4). Ten MIR169 genes are commonly up-regulated in all three HS conditions and can be considered as critical for thermo-tolerance (Figure 2c, d). Four MIR169s (Sly-MIR169a, a-2, b-3 and d-2) are uniquely up-regulated in harsh HS, suggesting significance in basal thermotolerance mechanisms. Sly-MIR169b-4 gene is up-regulated in acclimation and acquired HS but not in basal stress regime, suggesting its role in priming response of the plant for thermo-tolerance (Figure 2c, d). Ten MIR169 members continue to express at high levels even when the stress is removed during recovery phase (Figure 2c, d). These genes may be important components of HS memory, plant repair and healing machinery and good candidates for enhancing thermotolerance.

### MIR169s regulate tomato plant development

GUS aided promoter analysis in Arabidopsis has shown that most of the miR169s members show prominent expression in vascular tissues, miR169c expresses uniquely in shoot apex, root tip and lateral root primordia; mir169i/j and m/n express in the anther and pollen and miR169e exhibits maximum GUS activity in roots (Li et al. 2010; Zhao et al. 2011). To assess MIR169-mediated tissue-specific regulation in tomato, we did detailed investigation of MIR169 members in various tissues viz., leaf, root, inflorescence and fruit by qRT-PCR. We find that the MIR169 transcripts have low abundance at all stages of development (Supplementary Table S4) except for three loci (Sly-MIR169a, Sly-MIR169d-1, Sly-MIR169g). Keeping leaf as control, we find that MIR169s behave quite similarly in root and mature green (MG) fruit (Figure 2e, f). One striking difference is that Sly-MIR169b-1/b is highly abundant in root but is absent in MG and breaker fruits. Sly-MIR169d is not detected in roots, but is abundantly present in flower stages. Between the two inflorescence stages, Sly-MIR169a and a-1, are up-regulated in meiotic-stage only, while Sly-MIR169a-2 is down-regulated in both; Sly-MIR169b-1/b and ‘b-2/h’ are down-regulated in post-meiotic stages of flower development while Sly-MIR169e is oppositely regulated (Figure 2e, f). Sly-MIR169b-1/b is not detected in fruits and Sly-MIR169g is highly down-regulated (∼7-folds) in breaker stage. SlMIR169b-2 shows strong expression in roots and mature fruits, suggested its role in root and fruit development. Expression analysis of different MIR169 precursors highlights not only development-specific roles but also the regulation brought by MIR169 paralogs.

### MIR169 targets transcription factors and different classes of enzymes

Studies have shown that increase in the copy number of miRNA family is coupled with the increase in the number of targets (Maher et al. 2006; Sieber et al. 2007; Shi et al. 2016) thereby supporting the association of miRNA family functional diversification and expansion (Abrouk et al. 2012). Thus, enhanced MIR169 loci in tomato should also expand the targets it cleaves. To study this, a rigorous search was done using publicly available PARE libraries of tomato for miRNA169-mediated degradation of transcripts. We identified several targets for MIR169 in tomato (Supplementary Table S5). These include the classical targets of miR169s the transcription factor Nuclear Factor-Y A (NFY-A, five members) and several new targets. Several studies have also validated NY-FA family members, as the main targets of miR169 family members which are involved in the development of plant root architecture (Sorin et al. 2014), nodule formation (Zhao et al. 2011), disease resistance (Li et al. 2017), and stress responses (Zhao et al. 2009; Ni et al. 2013; Ding Zeng & He 2016; Luan et al. 2015). To validate the targets, we identified, we generated fusion constructs of targets with GFP reporter gene in its CDS or 3’ UTR, depending on the cleavage site of target. Cleavage of targets was confirmed by reduction in GFP transcripts levels by qRT-PCR when co-infiltrated with over-expression constructs of specific miR169 precursors in *N. benthamiana* leaves.

All five NF-YAs (NF-YA1, -A3, -A8, -A9, -A10) (Figure 3a) and seven novel targets were validated (Figure 3b). The novel targets include, other transcription factors like floral development gene SEPALLATA-3 (SEPL-3) and Auxin response factor-9B (ARF-9B); a chaperone DnaJ, enzymes like Aspartate aminotransferase (Asp-AT), Protein phosphatase 2C family protein (PP2C), RNA-binding-protein (RBP) and Tetratricopeptide-repeat-protein (TPR) that is important in protein-protein interactions (Figure 3b, Supplementary Figure S10). Overexpression of miR169a (miR169a-OX) reduces NF-YA3 and Asp-AT transcript by 2.4- and 3.3 folds, respectively. Very strong reduction is obtained for SEPL-3 (15.8 fold) and NF-YA9 (2.5 fold) by miR169b-OX overexpression. miR169c-OX overexpression causes reduction of NF-YA1 transcript by 2.5 fold. miR169d-OX mediates reduction of NF-YA8 (4.1 fold), NF-YA10 (4.4 fold) and ARF-9B (2.1 fold) transcripts. PP2C family protein transcripts are reduced (2.2 fold) by overexpression of miR169i (Figure 3a, b). To further confirm the miR169-mediated cleavage of all validated targets, miR169s expression was suppressed transiently by sponging-up mature miRNA by short-tandem-target-mimic (STTM) expressed under 2□×□35S promoter in tomato. Transcript levels of the target genes in STTM169 plants showed significant increased levels confirming them as true MIR169 targets (Figure 3c).

**Figure 3:**
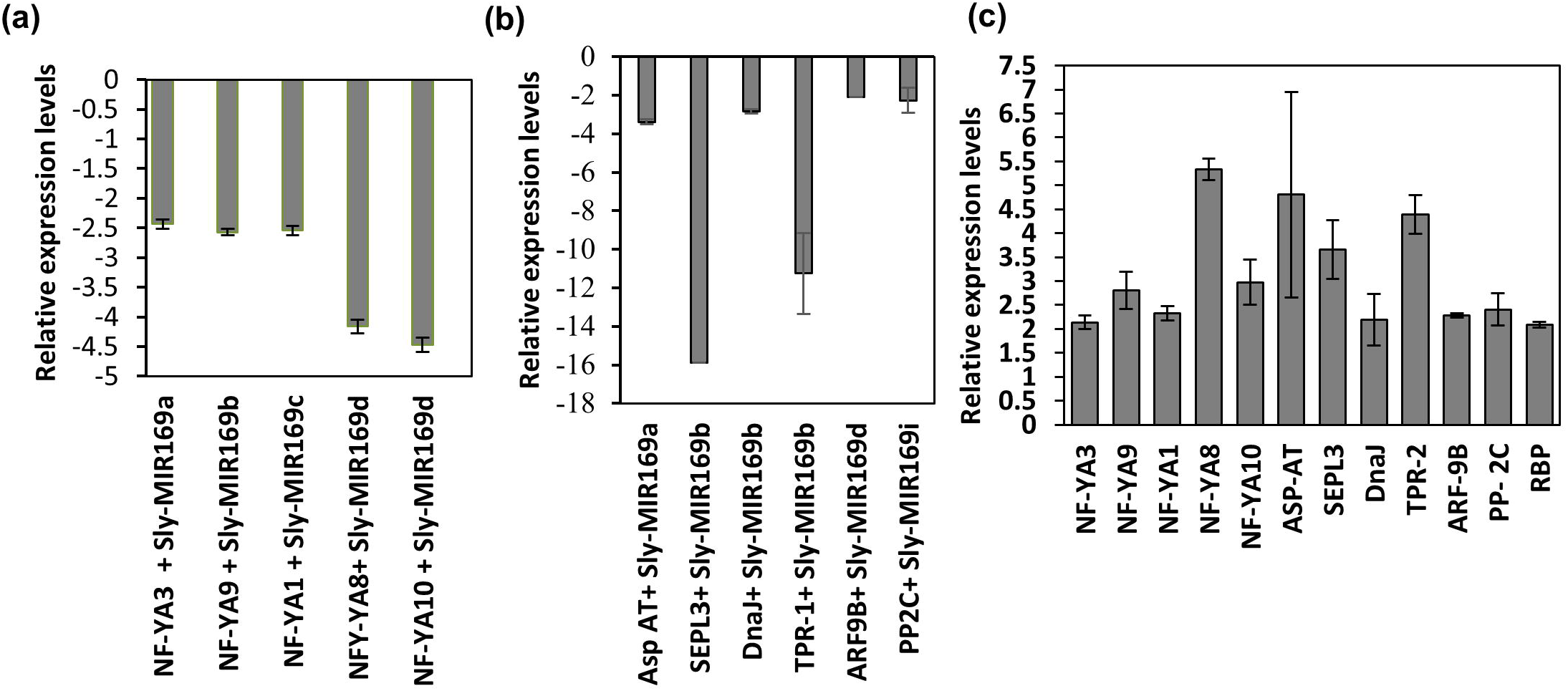
Target validation of MIR169 members by transient assays in *Nicotiana benthamiana* leaves. (a) Known targets (NF-YAs) of MIR169 members identified by degradome analysis and validated by transient assay using qRT-PCR. (b) qRT-PCR validation of novel targets of new MIR169 members identified by degradome analysis, ASP-AT - Aspartate aminotransferase, SEPL-3 - SEPALLATA-3, ARF-9b - Auxin response factor-9B, PP2C - Protein phosphatase 2C, TPR-2 - Tetratricopeptide repeat-2, DnaJ - Chaperone protein DnaJ. (c) qRT-PCR analysis of the accumulation of the target transcripts in STTM169 tomato lines. Data are presented as fold induction in STTMMIR169 infiltrated plant relative to STTM vector control plants. For both (a) and (b) relative expression levels of target were assessed by comparing the expression of the GFP fused to miR-sensitive targets alone and when co-infiltrated with different miR169 isoforms in *Nicotiana benthamiana* leaves. qRT-PCR analysis of GFP mRNAs of p35S: GFP-targets was performed with *hygromycin* gene that is present in the same vector as control for normalization. Data are shown as means ± Standard Deviation of three biological replicates with 3–4 agro-infiltrated plants for each construct.

To gain insights into probable functional prowess attained by MIR169 family because of new targets in tomato, we assessed the expression patterns of MIR:target pairs in different abiotic stresses and plant organs. Judging by the anti-correlation in expression patterns between miRNAs and targets, Sly-MIR169d:ARF-9B, Sly-MIR169i:PP2C and Sly-MIR169b-1:DnaJ modules are functional in dehydration stress. Regulation of DnaJ by miRNA169b is also apparent in cold and salt stress. miR169a mediated regulation of Asp-AT is significant in heat stress (Figure 4a-f). Zhenghua et al. (2019) have also reported the critical roles of PP2C in multiple abiotic stress responses. Previous studies in tomato have also revealed the involvement of ARFs in mediating auxin action in response to biotic and abiotic stresses (Bouzroud et al. 2018). Zhao et al. (2010) suggest that over-expression of DnaJ can confer NaCl-stress tolerance.

**Figure 4:**
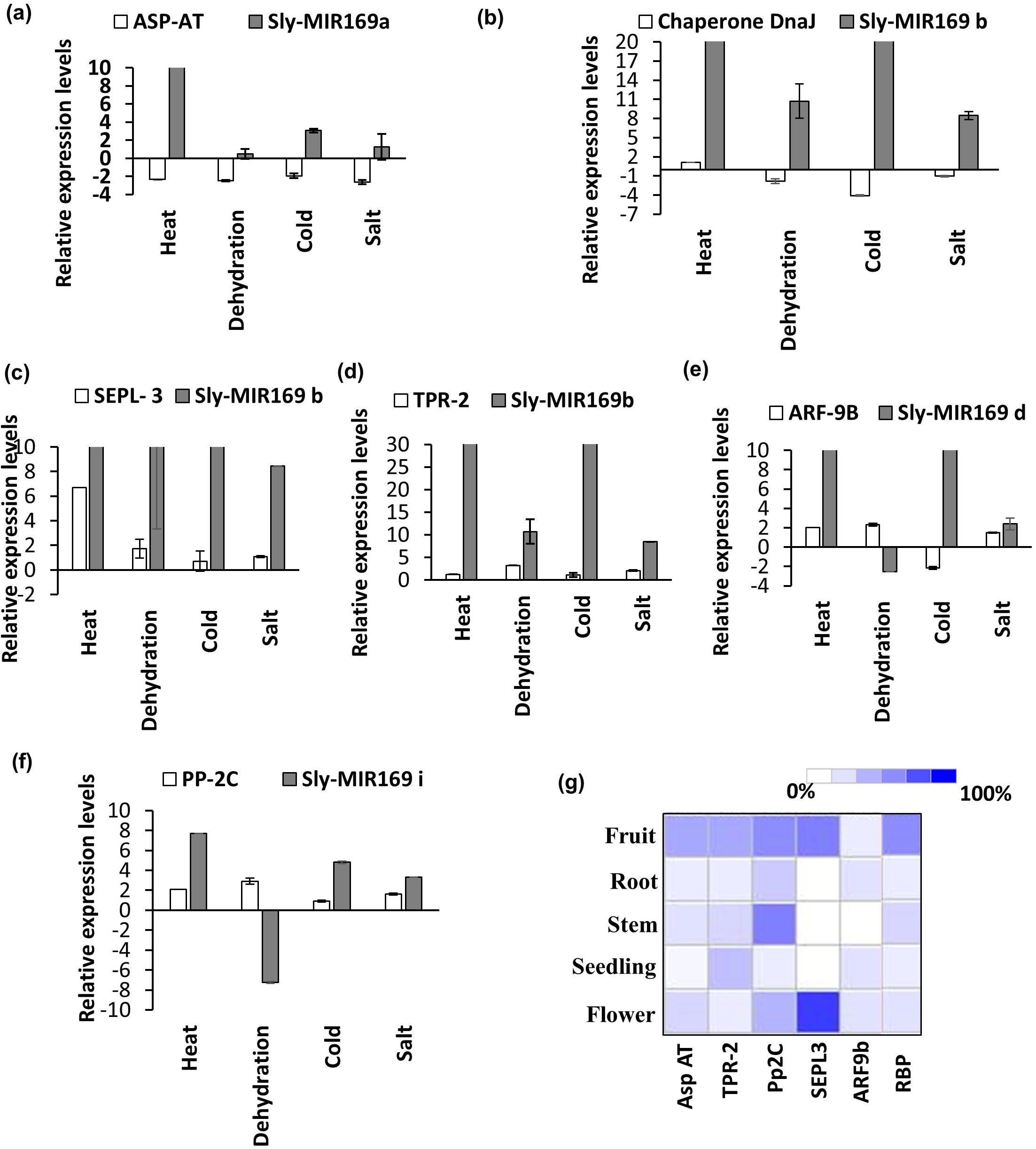
Establishing miRNA169: Novel target modules operating in different abiotic stresses and plant development. (a-f) Validation of miRNA169: novel target modules operating in abiotic stresses by comparing the expression profiles of MIR169s and targets. (g) Developmental expression patterns of the target genes obtained from Genevestigator. Results are given as heat maps in colored/white coding that reflect absolute signal values, where darker shade represents stronger expression. Targets and precursors were quantified by qRT-PCR. Data are shown as means ± SD of biological replicates and normalized to *Actin* reference gene using (2−ΔΔCT) method.

Furthermore, evaluation of targets expression in different organs of tomato by using microarray data available at Genevestigator revealed that miR169a-Asp-AT module is functional in flower and root. In literature, SEPL-3 regulates flower development (Pelaz et al. 2001; Castillejo et al. 2005); our analysis suggests its significance not only in inflorescence but also in roots and fruits (Figure 4g). Regulation of PP2C by mir169i is apparent in flower and root while miR169d regulates ARF-9B in flower and fruit (Figure 4g). Similar to our finding, functional analysis of a homolog of ARF-9B by DeJong et al. (2015) also indicates its roles in early tomato fruit development. It has been shown for TPR proteins that they play an important role in chloroplast gene expression (Hu et al. 2014), protein turnover (Park et al. 2007), photosystem assembly and repair (Park et al. 2007, Heinnickel et al. 2016), chlorophyll biosynthesis, and thylakoid membrane biogenesis (Schottkowski et al. 2009). We find that TPR activity that is regulated by miR169b is also important in root and fruit (Figure 4g). Our study shows that miR169h is present in only the cultivated variety so it was intriguing to establish role of any new target that may have been acquired during domestication. While six targets were predicted (Supplementary Table S4), we could validate only RBP as a bona fide target using STTM assay (Figure 3c). Inverse expression profiles of RBP and Sly-MIR169h were noted in tomato roots only (Figure 4g, 2e). The functional relevance of this needs further attention. Taking together our results and literature information confirms the significant roles of miR169:target modules in abiotic stresses and plant development (Supplementary Figure S11). The new targets thereby, enhance the regulatory diversity of this miRNA family.

### miR169 isoforms exhibit differential processing and regulate NF-YA10 in a spatial and stress-specific manner

It is known that secondary structural determinants of the precursor like stem-loops/bulges and sequences surrounding the miRNA/miRNA* region have important consequences on miRNA processing (Starega-Roslan et al. 2011 Mateos et al. 2010, Werner et al. 2010 Cuperus et al. 2010, Song et al. 2010 Bologna et al. 2013). The different miRNA169 precursors of tomato show variation in structures as well as differences in sequence in stem part of precursors (Supplementary Figure S3). We therefore, decided to evaluate, the processing efficacy of some of the known and the newly identified miRNA169 precursors. For assaying this, effector plasmids containing different MIR169 precursors under constitutive 35S promoter were co-infiltrated with a reporter plasmid containing the canonical MIR169 target NF-YA10 under the constitutive 35S promoter. As expected, NFY-A10 is cleaved by all MIR169 precursors however; there is great degree of variation in target reduction using different precursors. Strong reduction of NF-YA10 was mediated by Sly-MIR169b (6.5 fold down-regulation) followed by Sly-MIR169d-1, Sly-MIR169a, Sly-MIR169d, Sly-MIR169c and Sly-MIR169a-1, respectively (Figure 5a). Since all precursors are expressed under the same promoter therefore, post-transcriptional regulation of precursor processing itself and/or accumulation of mature miRNA is responsible for this differential cleavage of NF-YA10 transcripts.

**Figure 5:**
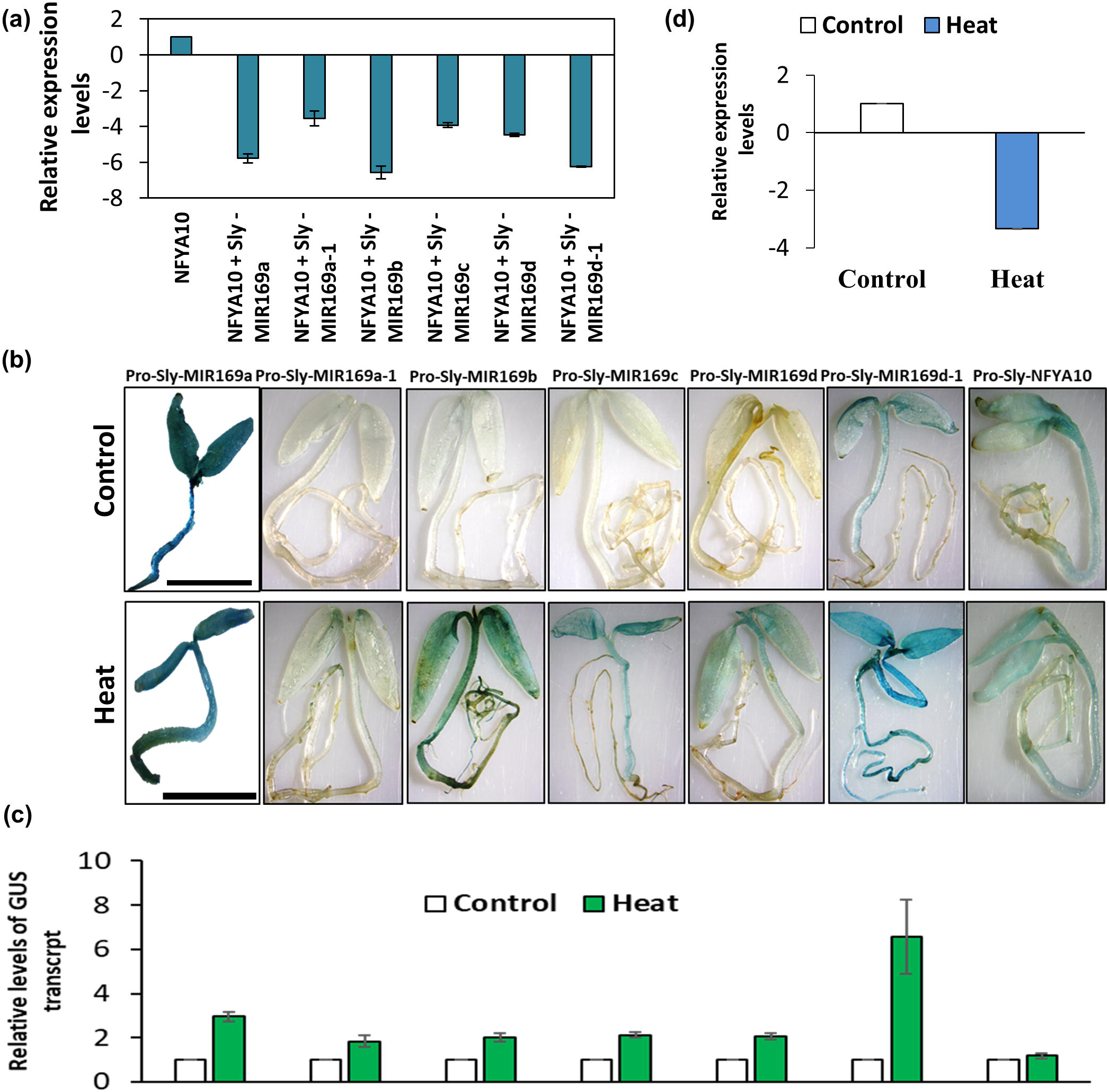
Differential processivity of MIR169 precursors regulates NF-YA10 cleavage level. (a) Cleavage of NF-YA10 by different MIR169 precursors as judged by qRT-PCR. Graph showing the transcript abundance of GFP:target transcript in absence and presence of miRNA precursor. Transcript levels were measured using GFP specific primers and *hygromycin* that is co-expressed in the same construct was used as normalization control to negate the infiltration variation. (b) Abundance of MIR169 precursors in different tissues and heat stress in comparison with NF-YA10 promoter activity. Each picture is representative of pool of seedlings used in the samples. (c) Levels of GUS transcripts measured by qRT-PCR after agro-infiltration of different MIR169s promoters:GUS reporter constructs in tomato seedlings. All qRT-PCR experiments were repeated at least three times on 70 seedlings per replicate with similar results, and average data is shown. Error bar denotes standard error. (d) Expression profile of NF-YA10 transcript in control and heat stress conditions.

Further, the question arises whether transcriptional control of precursor also regulates target abundance *in-planta*. Therefore, we dissected the interplay of the above MIR169s with NF-YA10 module at transcriptional regulation by comparing the ability of their promoters to drive GUS reporter gene in tomato seedlings. The promoters of Sly-MIR169a, Sly-MIR169d-1 and NF-YA10 express in entire seedling albeit weekly and appear to be the functional pairs under unstressed conditions in seedlings. HS induces the expression of all MIR169 loci in a tissue spatial manner, Sly-MIR169a, Sly-MIR169b and Sly-MIR169d-1 express constitutively while Sly-MIR169a-1, Sly-MIR169c and Sly-MIR169d express only in aerial above ground parts of tomato. Notably, while GUS transcripts do not increase in HS for the NF-YA10 promoter (Figure 5b, c), the mRNA abundance of NF-YA10 reduces by ∼4-folds (Figure 5d). This suggests only a post-transcriptional regulation of NF-YA10 by a combination of different MIR169 mediated cleavage during HS. Thus, NF-YA10 transcripts are regulated in a tissue-specific and stress-responsive manner by a combination of different MIR169 members (Figure 6b, c). Similarly, one can extrapolate that different MIR169 forms would be regulating different targets in spatio-temporal/other abiotic/biotic cues in tomato, thereby adding to regulatory complexity of this family.

### Conclusion

Plants are a unique group among eukaryotes that have a robust system to sense light and temperature to not only make their own food by photosynthesis but regulate other developmental switches including germination and flowering. Moreover, since they are rooted to one place, they are exposed to various biotic and abiotic stresses. Thus, for their successful survival, expanded gene families have evolved in plants that participate in networks of well-coordinated signaling pathways involving transcriptional, post-transcriptional and epigenetic controls. Micro RNAs regulate different aspects of plant development and response to external factors primarily by target cleavage and translational repression. MIR169 is an important evolutionary conserved miRNA family that has well established roles in root development, nodule regulation and is responsive to various biotic and abiotic stresses. In this work, we have identified eighteen functional loci in tomato that includes more precursors for MIR169a, b and d forms. The expansion of the family is a result of both tandem and segmental duplication during the course of evolution. Duplications have resulted in enhanced robustness by acquiring new promoters that are regulated in spatio-temporal as well as stress-responsive manner. Existence of polycistronic forms that are conserved between the cultivated and two wild varieties highlight an evolutionary conserved feature that has probably evolved for strong expression of miRNA it encodes. A specie-specific MIR169h form that is present only in the cultivated variety has also been validated. Moreover, we have shown differential cleavage efficiencies of the precursors that adds another layer of regulation that might be important for fine tuning the MIR169:target control mechanism in different stages of development and response to environment. We have identified four new mature forms that increase the possibility to engage new targets for functional divergence. In this regard, novel targets have been identified and validated that exhibit antagonist expression to miRNA in different abiotic stresses including dehydration, cold and salt stress. Also, the new targets appear to have roles in flower and fruit development also. This study increases the knowledge about MIR169 in tomato not only in terms of new members (both miRNA and targets) being identified but also their regulation, functional divergence and potential application for crop improvement.

## Supporting information

Supplementary Figure S1: Identification and validation of MIR169 family members in tomato

Supplementary Figure S2: Identification of duplicated MIR169 genes in tomato.

Supplementary Figure S3: Stem-loop structures of putative pre-miR169s in tomato.

Supplementary Figure S3: Stem-loop structures of putative pre-miR169s in tomato.

Supplementary Figure S4: Chromosomal localization, clustering and duplication of MIR169 family genes in tomato.

Supplementary Figure S5: Genomic organization of polycistronic clusters of miRNA169s on chromosome 7

Supplementary Figure S6: A species-specific MIR169h locus is present in cultivated tomato.

Supplementary figure S7: Comparison of MIR169b-1/b locus between wild and cultivated tomato species.

Supplementary Figure S8: Comparison of MIR169b2/h polycistronic locus between wild and cultivated tomato species.

Supplementary Figure S9: Acquisition of different promoters by MIR169 paralog pairs in tomato.

Supplementary Figure S10: Alignment of novel targets with miR169 in tomato.

Supplementary Figure S11: The miRNA169: target functional module on the basis of pathway analysis in tomato.

Supplementary Table S1: List of primers used in the study.

Supplementary Table S2: Comparison of MIR169 loci in wild and cultivated tomato species.

Supplementary Table S3: Promoter analysis of Sly-MIR169 family members in tomato.

Supplementary Table S4: qRT-PCR based expression data of Sly-MIR169 members in different abiotic stresses and plant organs.

Supplementary Table S5: Targets of Sly-MIR169 members identified by degradome analysis.

## Acknowledgments

The authors acknowledge NIPGR core grant, phytotron facility, CIF and field area. CLN1621L seeds were provided by AVRDC, TAIWAN. SR and SJ acknowledge Department of Biotechnology (DBT) and University Grants Commission (UGC) Govt. of India, respectively for the award of research fellowships.

## Supplementary Figure legends

**Supplementary Figure S1: Identification and validation of MIR169 family members in tomato. (**a) Sequence homology based bioinformatics pipeline for identification of MIR69 genes in tomato. (b) Tabulated total MIR169 family members of tomato (c) Sequence alignment of 9 mature forms of miR169s in tomato. For precursor identification a repertoire of miR169 sequences from miRBASE were used as query in BLASTn on tomato chromosome build.v.4 available at Sol Genomics Network. For validation of predicted precursors, cDNA was synthesized from pooled RNA from control and heat stress leaf tissue. Precursors were amplified and cloned into TA vector and confirmed by sequencing. At least two clones were sequenced for individual precursor.

**Supplementary Figure S2: Identification of duplicated MIR169 genes in tomato. (**a) Sequence alignments of precursor sequences of paralogs with high degree of sequence similarity, percent similarity between paralogs is indicated. The nucleotide mismatches are colored light blue and reflected as gaps in ‘consensus’ bar. Any indel is reflected as a gap in the ‘occupancy’ bar. (b) Phylogenetic analysis of MIR169 members to identify pairs of duplicated genes, depicted by red boxes. The unrooted tree was constructed with precursor sequences by neighbor joining algorithm using clustalx software. Reliability values at each branch represent bootstrap samples (1000 replicates). The scale bar represents the nucleotide substitution per site.

**Supplementary Figure S3: Stem-loop structures of putative pre-miR169s in tomato.** The mature miRNA/miRNA* portion is highlighted in black boxes. The secondary structure of the precursors of the identified miRNAs was predicted using the RNAfold web server and to designate new miRNAs, we adopted the criteria proposed previously (Zhang et al. 2006a; Meyers et al. 2008).

**Supplementary Figure S4: Chromosomal localization, clustering and duplication of MIR169 family genes in tomato**. Each box represents a chromosome numbered on top. The approximate position of each tomato MIR169 gene is marked on the chromosome with a short black line. MIR169 family comprises of four gene clusters depicted by pink bars. Duplicated genes are indicated with red brackets (tandem) and green brackets (segmental). Duplication of genes was determined by location, phylogeny and sequence identity.

**Supplementary Figure S5: Genomic organization of polycistronic clusters of miRNA169s on chromosome 7**. (a) Pictorial depiction of MIR169d to h cluster on chromosome 7. Mature miRNA sequences are indicated with a small colored line in precursor shown as hairpin-loop structure, promoter regions and transcriptional orientations are indicated with red arrows. Approximate position of primers used for confirming polycistronic nature are depicted with green arrows. Distances between precursors has been marked by open-ended arrows. (b) Putative polycistronic pair secondary structure predicted by RNA fold software. The mature miRNA/miRNA* portion is highlighted in black boxes. (c) PCR amplicons of MIR169b-1/b and MIR169b-2/h using genomic DNA and cDNA using primers as shown in (a) to validate polycistronic nature.

**Supplementary Figure S6: A species-specific MIR169h locus is present in cultivated tomato**. (a) MIR169b2/h locus amplified using genomic DNA (gDNA) and cDNA of cultivated (*Solanum lycopersicum)* and wild species (*S. pimpinellifolium)*. Amplicons were further verified by sequencing. M: DNA marker, S. lyco: *Solanum lycopersicum* and S. pimp: *Solanum pimpinellifolium*. (b) Sequence comparison of MIR169b-2/h locus from wild and cultivated species to confirm deletion and sequence rearrangements in precursor and mature miR169h. Mature miRNA sequences are highlighted by yellow color. Red lines in between the aligned sequences depict the deleted region of *Solanum lycopersicum* MIR169b2/h locus. Vertical red colored box represents the changed base in *S. lycopersicum* that leads to generation of miR169h.

**Supplementary figure S7: Comparison of MIR169b-1/b locus between wild and cultivated tomato species.** Sequence comparison of MIR169b-1/b locus from wild and cultivated species to confirm deletion and sequence arrangements in precursor and mature miR169b. Red and black colored boxes represent MIR169b-1 and MIR169b precursors, respectively. The red dotted line depicts deleted region in *S. pennellii* of the inter-precursor region of MIR169b-1/b polycistronic locus.

**Supplementary Figure S8: Comparison of MIR169b2/h polycistronic locus between wild and cultivated tomato species.** Sequence comparison of MIR169b-2/h locus from wild and cultivated species to confirm deletion and sequence arrangements in precursor and mature miR169. Red and black colored boxes represent MIR169b-2 and MIR169h precursors, respectively. The red dotted line depicts deleted region in S. pennellii of the inter-precursor region of MIR169b-2/h polycistronic locus.

**Supplementary Figure S9: Acquisition of different promoters by MIR169 paralog pairs in tomato.** (a-c) GUS aided promoter activity analysis of paralog pairs Sly-MIR169a and a-2, Sly-MIR169b-3 and b-4 and Sly-MIR169b-1/b and Sly-MIR169b2/h in tomato seedlings. For GUS reporter assay, 2 kb genomic sequences upstream of the precursor start sites were used. Staining was performed 2 days after agro-infiltration in 4-days-old tomato seedlings. These experiments were repeated at least three times with 70 seedlings per replicate with similar results; a representative picture from each experiment is shown.

**Supplementary Figure S10: Alignment of novel targets with miR169 in tomato**. The alignments of fragments of target mRNAs that have complementarity to miR169s are shown. Alignments of miRNA and targets were obtained from psRNATARGET prediction server.

**Supplementary Figure S11: The miRNA169: target functional module on the basis of pathway analysis in tomato**. Pathway analysis of MIR169 novel targets reveals their involvement in different biological processes including, signalling, stress tolerance, flowering and metabolic pathways.

## Supplementary Table legends

**Supplementary Table S1: List of primers used in the study.**

**Supplementary Table S2: Comparison of MIR169 loci in wild and cultivated tomato species.**

**Supplementary Table S3: Promoter analysis of Sly-MIR169 family members in tomato.**

**Supplementary Table S4: qRT-PCR based expression data of Sly-MIR169 members in different abiotic stresses and plant organs.**

**Supplementary Table S5: Targets of Sly-MIR169 members identified by degradome analysis.**

## Literature cited

Abrouk M, Zhang R, Murat F, Li A, Pont C, Mao L and Salse J (2012) Grass microRNA gene paleo history unveils new insights into gene dosage balance in sub-genome partitioning after whole-genome duplication. The Plant Cell, 24:1776–1792.

Altuvia Y, Landgraf P, Lithwick G, Elefant N (2005) Clustering and conservation patterns of human microRNAs. Nucleic Acids Research, 33:2697–706.

Axtell MJ, Bartel DP (2005) Antiquity of microRNAs and their targets in land plants. The Plant Cell, 17:1658–1673.

Bai Q, Wang X, Chen X, Shi G, Liu Z, Guo C and Xiao K (2018) Wheat miRNA TaemiR408 acts as an essential mediator in plant tolerance to Pi deprivation and salt stress via modulating stress-associated physiological processes. Frontier in plant Sciences, doi:10.3389/fpls.2018.00499. eCollection 2018.

Baker CC, Sieber P, Wellmer F, Meyerowitz EM (2005) The early extra petals1 mutant uncovers a role for microRNA miR164c in regulating petal number in Arabidopsis. Current Biology, 15:303–315.

Bartel, D.P. (2004) MicroRNAs: Genomics, biogenesis, mechanism, and function. Cell 116:281–297

Bhan B, Koul A, Sharma D, Manzoor MM, Kaul S, Gupta S, Dhar MK (2019) Identification and expression profiling of miRNAs in two color variants of carrot (Daucus carota L.) using deep sequencing. PLOS ONE, doi.org/10.1371/journal.pone.0212746.

Bologna NG, Schapire AL, Zhai J, Chorostecki U, Boisbouvier J, Meyers BC, Palatnik JF (2013) Multiple RNA recognition patterns during microRNA biogenesis in plants. Genome Research, 23: 1675–1689.

Bouzroud S, Gouiaa S, Hu N, Bernadac A, Mila I, Bendaou N, Smouni A, Bouzayen M, Zouine M (2018) Auxin Response Factors (ARFs) are potential mediators of auxin action in tomato response to biotic and abiotic stress (Solanum lycopersicum). PLOS ONE, doi.org/10.1371/journal.pone.0193517.

Buhtz A, Pieritz J, Springer F, Kehr J (2010) Phloem small RNAs, nutrient stress responses, and systemic mobility. BMC Plant Biology, doi: 10.1186/1471-2229-10-64.

Calviño M, Messing J (2013) Discovery of MicroRNA169 Gene Copies in Genomes of Flowering Plants through Positional Information. Genome Biology and Evolution, 5:402–417.

Cannon SB, Mitra A, Baumgarten A, Young ND, May G (2004) The roles of segmental and tandem gene duplication in the evolution of large gene families in *Arabidopsis thaliana*. BMC Plant Biology, doi: 10.1186/1471-2229-4-10.

Castillejo C, Romera-Branchat M, Pelaz S (2005) A new role of the Arabidopsis SEPALLATA3 gene revealed by its constitutive expression. Plant Journal, 43:586–596.

Chorostecki U, Moro B, Rojas A M L, Debernardi J M, Schapire A L, Notredame C and Palatnik J F (2017) Evolutionary Footprints Reveal Insights into Plant MicroRNA Biogenesis, The Plant Cell, 29:1248–1261.

Combier JP, Frugier F, de Billy F, Boualem A, El-Yahyaoui F, Moreau S, Vernié T, Ott T, Gamas P, Crespi M, Niebel A (2006) MtHAP2-1 is a key transcriptional regulator of symbiotic nodule development regulated by microRNA169 in Medicago truncatula. Genes and Development 5:3084–8.

Cuperus JT, Carbonell A, Fahlgren N, Garcia-Ruiz H, Burke RT, Takeda A, Sullivan CM, Gilbert SD, Montgomery TA, Carrington JC (2010) Unique functionality of 22-nt miRNAs in triggering RDR6-dependent siRNA biogenesis from target transcripts in Arabidopsis. Nature Structural Molecular Biology, 17:997–1003.

de Jong M, Wolters-Arts M, Schimmel BC, Stultiens CL, de Groot PF, Powers SJ, Tikunov YM, Bovy AG, Mariani C, Vriezen WH, Rieu I (2015) Solanum lycopersicum AUXIN RESPONSE FACTOR 9 regulates cell division activity during early tomato fruit development. Journal of Experimental Botany, 11:3405–16.

Ding Q, Zeng J, and He XQ (2016) MiR169 and its target PagHAP2-6 regulated by ABA are involved in poplar cambium dormancy. Journal of plant physiology, 198:1–9.

Du Q, Zhao M, Gao W, Sun S and Li W (2017) microRNA/microRNA* complementarity is important for the regulation pattern of NFYA5 by miR169 under dehydration shock in Arabidopsis. Plant Journal, 91: 22–33.

Guan Q, Lu X, Zeng H, Zhang Y, Zhu J (2013) Heat stress induction of miR398 triggers a regulatory loop that is critical for thermotolerance in Arabidopsis. The Plant Journal, 74: 840–851.

Guddeti S, Zhang DC, Li AL, Leseberg CH, Kang H, Li XG, Zhai WX, Johns MA, Mao L (2005) Molecular evolution of the rice miR395 gene family. Cell Research, 15:631–638.

He Z, Wu J, Sun X, Dai M (2019) The Maize Clade A PP2C Phosphatases Play Critical Roles in Multiple Abiotic Stress Responses. International Journal of Molecular Sciences, doi: 10.3390/ijms20143573.

Heinnickel M, Kim R, Wittkopp T, Yang W, Walters K, Herbert S (2016) Tetratricopeptide repeat protein protects photosystem I from oxidative disruption during assembly. Proceedings of the National Academy of Sciences, 113:10.1073/pnas.1524040113.

Hivrale V, Zheng Y, Puli COR, Jagadeeswaran G, Gowdu K, Kakani VG, Barakat A, Sunkar R (2016) Characterization of drought□ and heat□responsive microRNAs in switchgrass. Plant Science, 242: 214–223.

Hu Z, Xu F, Guan L, Qian P, Liu Y, Zhang H, Huang Y, Hou S (2014) The tetratricopeptide repeat-containing protein slow green1 is required for chloroplast development in Arabidopsis. Journal of Experimental Botany, 4:1111–23.

Hua W, Hua G, Han B (2009) Genome-wide survey and expression profiling of heat shock proteins and heat shock factors revealed overlapped and stress specific response under abiotic stresses in rice. Plant Science, 176:583–590.

Jagtap S, and Shivaprasad P V (2014) Diversity, expression and mRNA targeting abilities of Argonaute-targeting miRNAs among selected vascular plants. BMC Genomics, doi:10.1186/1471-2164-15-1049

Kansal S, Mutum RD, Balyan SC, Arora MK, Singh AK, Mathur S, Raghuvanshi S (2015) Unique miRNome during anthesis in drought-tolerant indica rice var. Nagina 22. Planta, 6:1543–59.

Kulcheski FR, Luiz FV de O, Molina LG, Almerão MP, Rodrigues FA, Marcolino J, Barbosa JF, Moreira RS, Nepomuceno AL, Francismar C Marcelino-Guimarães, Abdelnoor RV, Nascimento LC, Carazzolle MF, Gonçalo AG Pereira & Rogério Margis (2011) Identification of novel soybean microRNAs involved in abiotic and biotic stresses. BMC Genomics, doi:10.1186/1471-2164-12-307.

Lee H, Yoo SJ, Lee JH, Kim W, Yoo SK, Fitzgerald H, Carrington JC, Ahn JH (2010) Genetic framework for flowering□time regulation by ambient temperature□responsive miRNAs in Arabidopsis. Nucleic acids research, 38: 3081–3093.

Li S, Li K, Ju Z, Cao D, Fu D, Zhu H, Zhu B, Luo Y (2016) Genome-wide analysis of tomato NF-Y factors and their role in fruit ripening. BMC Genomics, doi:10.1186/s12864-015-2334-2.

Li WX, Oono Y, Zhu J, He XJ, Wu JM, Iida K, Lu XY, Cui X, Jin H, Zhu JK (2008) The Arabidopsis NFYA5 transcription factor is regulated transcriptionally and post-transcriptionally to promote drought resistance. The Plant Cell, 20:2238–2251.

Li X, Xie X, Li JC, Hou Y, Zhai L, Wang X, Fu Y, Liu R, Bian S (2017) Conservation and diversification of the miR166 family in soybean and potential roles of newly identified miR166s. BMC Plant Biology, doi:10.1186/s12870-017-0983-9.

Li Y, Fu Y, Ji L, Wu C-A, Zheng C (2010) Characterization and expression analysis of the Arabidopsis mir169 family. Plant Science, 178:271–280.

Li Y, Zhao SL, Li JL, Hu XH, Wang H, Cao XL, Xu YJ, Zhao ZX, Xiao ZY, Yang N, Fan J, Huang F, Wang WM (2017) Osa-miR169 negatively regulates rice immunity against the blast fungus Magnaportheoryzae. Frontiers in Plant Science, doi: 10.3389/fpls.2017.00002.

Liang G, He H and Yu D (2012) Identification of nitrogen starvation responsive microRNAs in Arabidopsis thaliana. PLoS One, doi: 10.1371/journal.pone.0048951.

Lu Y and Yang X (2010) Computational Identification of Novel MicroRNAs and Their Targets in Vigna unguiculata. Comparative and functional genomics, 1531-6912:F1220–36.

Luan M, Xu M, Lu Y, Zhang L, Fan Y, and Wang L (2015) Expression of zma-miR169 miRNAs and their target ZmNF-YA genes in response to abiotic stress in maize leaves. Gene, 555:178–185.

Maher C, Stein L, Ware D (2006) Evolution of Arabidopsis microRNA families through duplication events. Genome Research, 16:510–519.

Maher C, Stein L, Ware D. (2006) Evolution of Arabidopsis microRNA families through duplication events. Genome Research, 16:510–519.

Marcinkowska M, Szymanski M, Krzyzosiak WJ, Kozlowski P (2011) Copy number variation of microRNA genes in the human genome. BMC genomics, doi: 10.1186/1471-2164-12-183.

Mateos JL, Bologna NG, Chorostecki U, Palatnik JF (2009) Identification of microRNA processing determinants by random mutagenesis of Arabidopsis MIR172a precursor. Current Biology, 1:49–54.

Merchan F, Boualem A, Crespi M, Frugier F (2009) Plant poly-cistronic precursors containing non-homologous microRNAs target transcripts encoding functionally related proteins. Genome Biology, 10:R136

Mi S, Cai T, Hu Y, Chen Y, Hodges E, Ni F, Wu L, Li S, Zhou H, Long C, Chen S, Hannon GJ, Qi YJ (2008) Sorting of Small RNAs into Arabidopsis Argonaute complexes is directed by the 5′ terminal nucleotide, Cell,1:116–127.

Mica E, Gianfranceschi L, Pè ME (2006) Characterization of five microRNA families in maize. Journal of Experimental Botany, 57:2601–12.

Muoki RC, Paul A, Kumari A, Singh K, Kumar S (2012b) An improved protocol for the isolation of RNA from roots of tea (*Camellia sinensis* (L.) *O*. Kuntze). Molecular Biotechnology, 52:82–88.

Mutum, R D, Kumar S, Balyan S, Kansal S, Mathur S, & Raghuvanshi S (2016). Identification of novel miRNAs from drought tolerant rice variety Nagina 22. Scientific reports, 6:30786.

Ni Z, Hu Z, Jiang Q, and Zhang H (2013) GmNFYA3, a target gene of miR169, is a positive regulator of plant tolerance to drought stress. Plant molecular biology, 82:113–129.

Pan Y, Niu M, Liang J, Lin E, Tong Z, Zhang J (2017) identification of heat-responsive miRNAs to reveal the miRNA-mediated regulatory network of heat stress response in Betulaluminifera. Trees, 31:1635–1652.

Pant BD, Musialak□Lange M, Nuc P, May P, Buhtz A, Kehr J, Walther D, Scheible WR (2009) Identification of nutrient□responsive Arabidopsis and rapeseed microRNAs by comprehensive real□time polymerase chain reaction profiling and small RNA sequencing. Plant physiology, 150:1541–1555.

Park S, Khamai P, Garcia-Cerdan JG, Melis A (2007) REP27, a tetratricopeptide repeat nuclear-encoded and chloroplast-localized protein, functions in D1/32-kD reaction center protein turnover and photosystem II repair from photodamage. Plant Physiology, 143:1547–1560.

Paul A, Rao S, Mathur S (2016) The α-Crystallin Domain Containing Genes: Identification, Phylogeny and Expression Profiling in Abiotic Stress, Phytohormone Response and Development in Tomato (Solanum lycopersicum). Frontiers in Plant Sciences, doi: 10.3389/fpls.2016.00426.

Pelaz S, Tapia-López R, Alvarez-Buylla ER, Yanofsky MF (2001) Conversion of leaves into petals in Arabidopsis. Current Biology, 3:182–184.

Reinhart BJ, Weinstein EG, Rhoades MW, Bartel B, Bartel DP (2002) MicroRNAs in plants. Genes and Development, 16:1616–1626.

Salvador-Guirao R, Hsing YI, & San Segundo B (2018) The Polycistronic miR166k-166h Positively Regulates Rice Immunity via Post-transcriptional Control of EIN2. Frontiers in plant science, doi: 10.3389/fpls.2018.00337.

Sarkar K N, Kim YK, Grover A (2009) Rice sHsp genes: genomic organization and expression profiling under stress and development. BMC genomics, doi: 10.1186/1471-2164-10-393.

Schottkowski M, Ratke J, Oster U, Nowaczyk M, Nickelsen J (2009) Pitt, a novel tetratricopeptide repeat protein involved in light-dependent chlorophyll biosynthesis and thylakoid membrane biogenesis in Synechocystis sp. PCC 6803. Molecular Plant, 2:1289–1297.

Sieber P, Wellmer F, Gheyselinck J, Riechmann JL, Meyerowitz EM (2007) Redundancy and specialization among plant microRNAs: role of the MIR164 family in developmental robustness, Development, 134:1051–1060.

Song L, Axtell MJ, Fedoroff NV (2010) RNA secondary structural determinants of miRNA precursor processing in Arabidopsis, Current Biology, 20:37–41.

Song S, Xu Y, Huang D, Ashraf MA, Li J, Hu W, Jin Z, Zeng C, Tang F, Xu B, Zeng H, Li Y, Xie J (2018) Identification and characterization of miRNA169 family members in banana (*Musa acuminata* L.) that respond to fusarium oxysporum f. sp. cubense infection in banana cultivars. Peer Journal, doi: 10.7717/peerj.6209.

Sorin C, Declerck M, Christ A, Blein T, Ma L, Lelandais□Brière C, Njo MF, Beeckman T,Crespi M, and Hartmann C (2014) A miR169 isoform regulates specific NF□YA targets and root architecture in Arabidopsis. New Phytologist, 202:1197–1211.

Starega-Roslan J, Koscianska E, Kozlowski P, Krzyzosiak W J (2011) The role of the precursor structure in the biogenesis of microRNA. Cellular and Molecular Life Sciences, 68:2859–2871.

Sunkar R, Jagadeeswaran G (2008) In silico identification of conserved microRNAs in large number of diverse plant species. BMC Plant Biology, doi: 10.1186/1471-2229-8-37.

Tanzer A and Stadler PF (2004) Molecular Evolution of a MicroRNA Cluster. Journal of Molecular Biology, 339:327–335.

Thakur V, Wanchana S, Xu M, Bruskiewich R, Quick WP, Mosig A, Zhu XG. (2011) Characterization of statistical features for plant microRNA prediction. BMC genomics, doi: 10.1186/1471-2164-12-108.

Thiebaut F, Rojas C A, Grativol C, Motta M R, Vieira T, Regulski M (2014) Genome-wide identification of microRNA and siRNA responsive to endophytic beneficial diazotrophic bacteria in maize. BMC genomics, doi: 10.1186/1471-2164-15-766.

Voorrips R E (2002) MapChart: software for the graphical presentation of linkage maps and QTLs. Journal of heredity, 93:77–78.

Xu F, Liu Q, Chen L, Kuang J, Walk T, Wang J, Liao H (2013) Genome wide identification of soybean microRNAs and their targets reveals their organ-specificity and responses to phosphate starvation. BMC genomics, doi: 10.1186/1471-2164-14-66.

Xu MY, Zhang L, Li WW, Hu XL, Wang MB, Fan YL, Zhang CY, Wang L (2014) Stress-induced early flowering is mediated by miR169 in Arabidopsis thaliana. Journal of Experimental Botany 65(1):89–101.

Yu C, Chen Y, Cao Y, Chen H, Wang J, Bi Y-M, Tian F, Yang F, Rothstein SJ, Zhou X (2018) Overexpression of miR169o, an overlapping microRNA in response to both nitrogen limitation and bacterial infection, promotes nitrogen use efficiency and susceptibility to bacterial blight in rice. Plant and Cell Physiology, 59:1234–1247.

Zhang BH, Pan XP, Cox SB, Cobb GP, Anderson TA (2006) Evidence that miRNAs are different from other RNAs. Cellular and Molecular Life Sciences, 63:246–54.

Zhang X, Zou Z, Gong P, Zhang J, Ziaf K, Li H, Xiao F, and Ye Z (2011) Over-expression of microRNA169 confers enhanced drought tolerance to tomato. Biotechnology letters, 33:403–409.

Zhao B, Ge L, Liang R, Li W, Ruan K, Lin H, and Jin Y (2009) Members of miR-169 family are induced by high salinity and transiently inhibit the NF-YA transcription factor. BMC molecular biology, doi: 10.1186/1471-2199-10-29.

Zhao B, Liang R, Ge L, Li W, Xiao H, Lin H, Ruan K, Jin Y (2007) Identification of drought-induced microRNAs in rice. Biochemical and Biophysical Research Communications, 354:585–590.

Zhao M, Ding H, Zhu JK, Zhang F, Li WX (2011) Involvement of miR169 in the nitrogen-starvation responses in Arabidopsis. New Phytologist, 190:906–1.

Zhao M, Tai H, Sun S, Zhang F, Xu Y and Li WX (2012) Cloning and characterization of maize miRNAs involved in responses to nitrogen deficiency. PLoS One, doi: 10.1371/journal.pone.0029669.

Zhao Z, Zhang W, Yan J, Zhang J, Liu Z L X and Yi Yang (2010) Over-expression of Arabidopsis DnaJ (Hsp40) contributes to NaCl-stress tolerance. African Journal of Biotechnology, doi: 10.5897/AJB09.1450.

Zhou X, Wang G, Sutoh K, Zhu JK, Zhang W (2008) Identification of cold□inducible microRNAs in plants by transcriptome analysis. Biochim et Biophys Acta,11:780–8.

Zhu H, Zhang Y, Tang R, Qu H, Duan X, Jiang Y (2019) Banana sRNAome and degradome identify microRNAs functioning in differential responses to temperature stress. BMC genomics, doi: 10.1186/s12864-018-5395-1.

Zuker M (2003) M fold web server for nucleic acid folding and hybridization prediction. Nucleic Acids Research, 13:3406–3415.

