## Supplementary figures and images for "Novel insights into expansion and functional diversification of MIR169 family in tomato"

### Supplementary Figure S1: Identification and validation of MIR169 family members in tomato

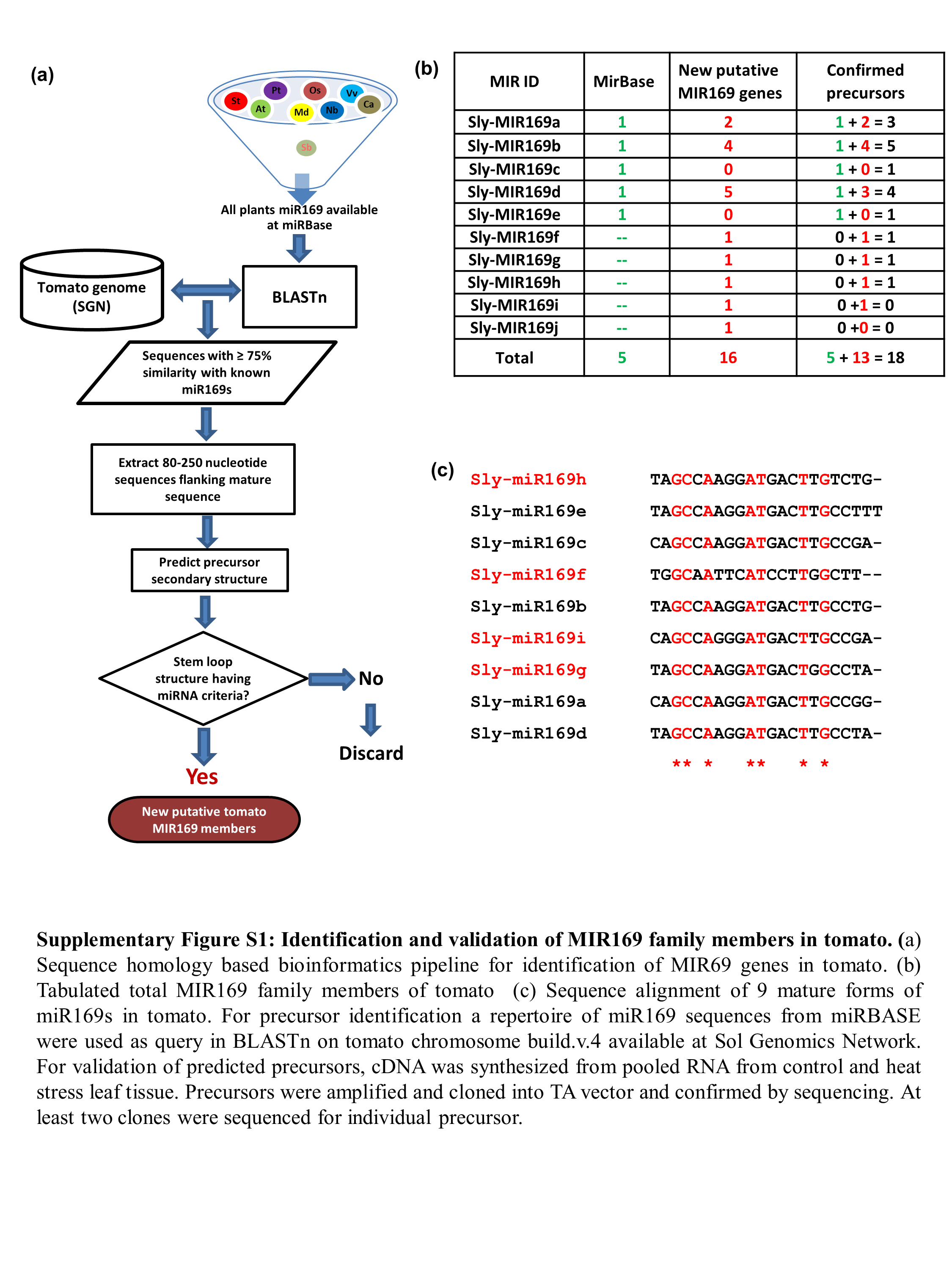

### Supplementary Figure S2: Identification of duplicated MIR169 genes in tomato.

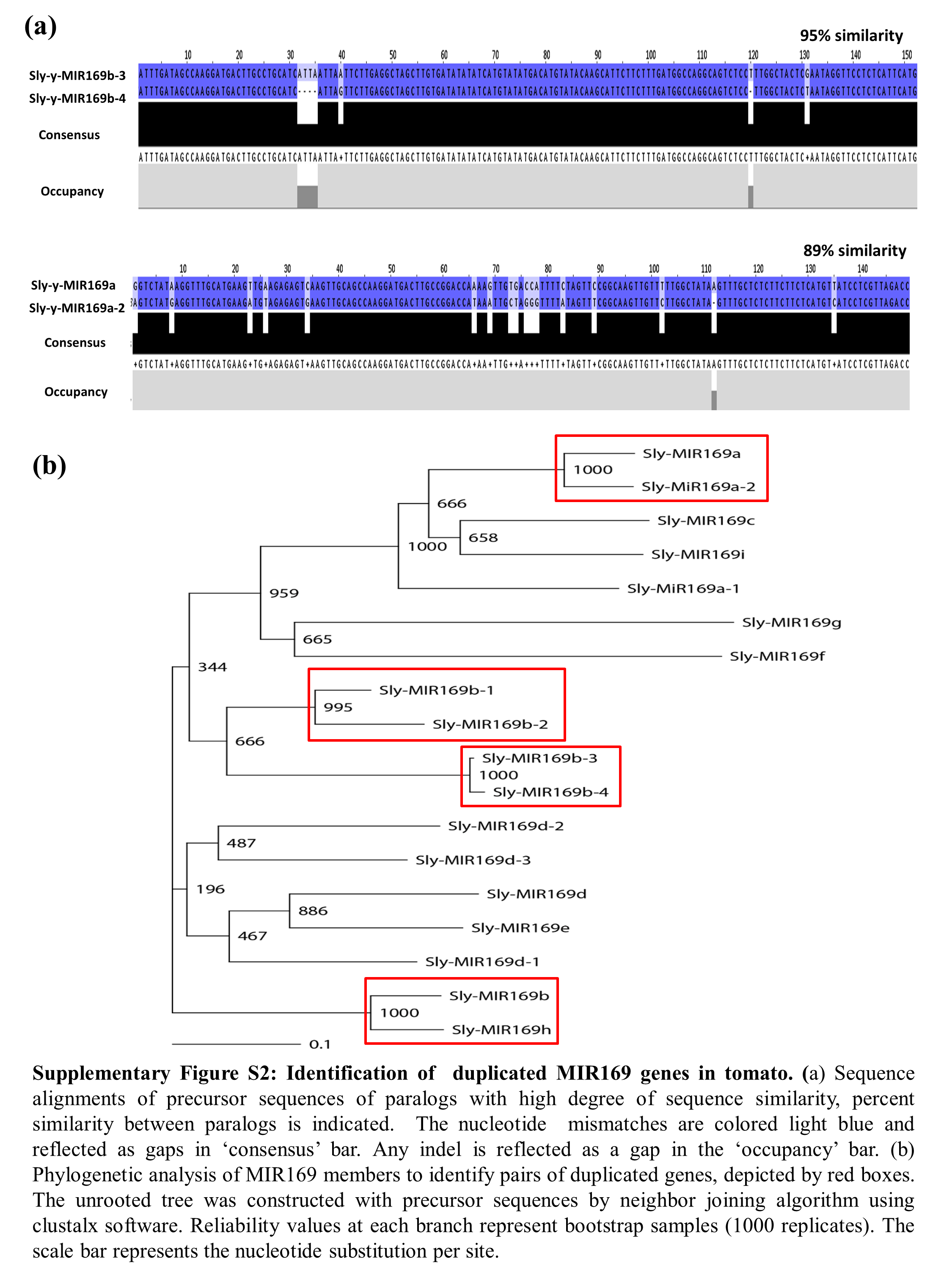

### Supplementary Figure S3: Stem-loop structures of putative pre-miR169s in tomato.

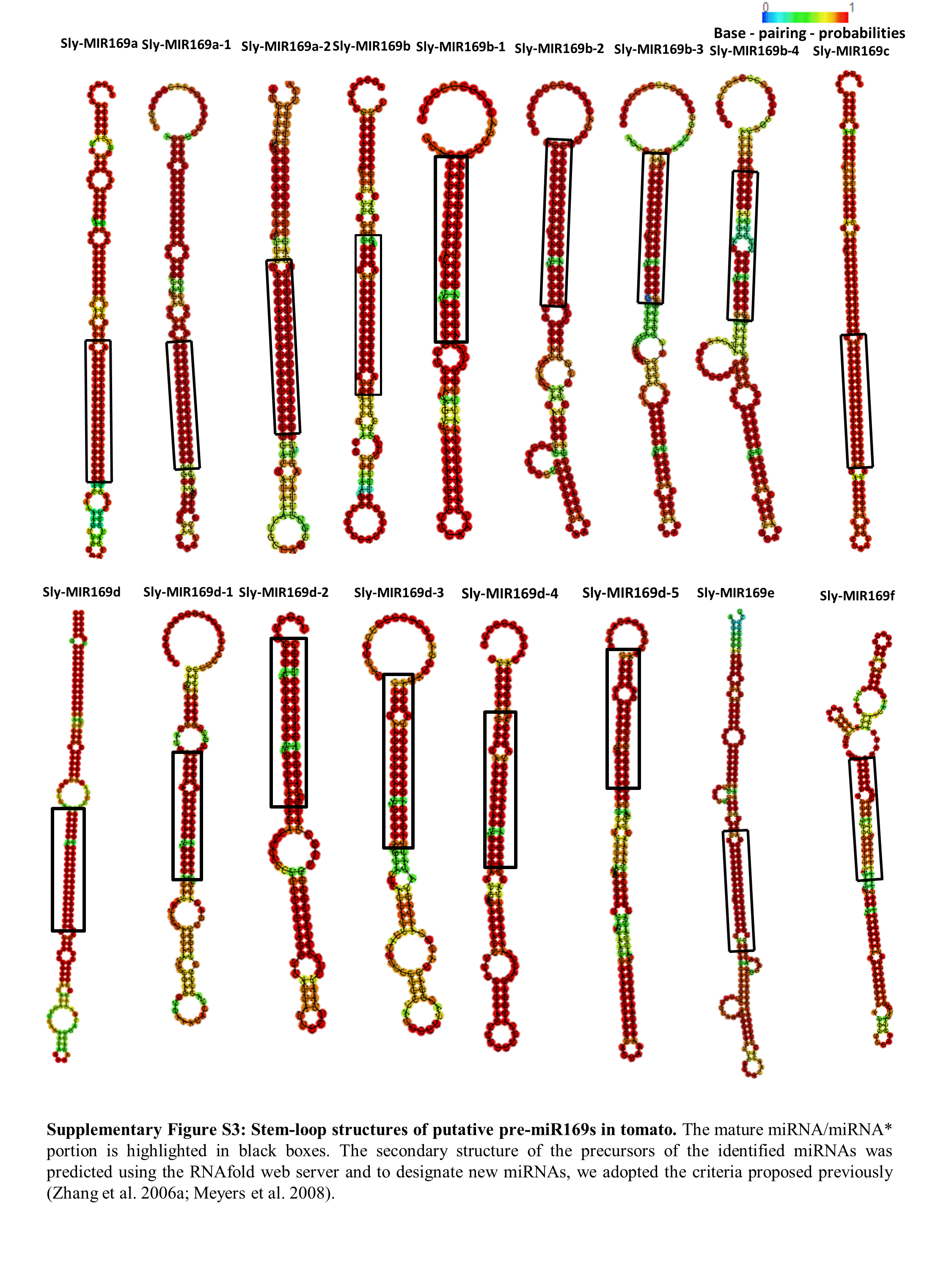

### Supplementary Figure S3: Stem-loop structures of putative pre-miR169s in tomato.

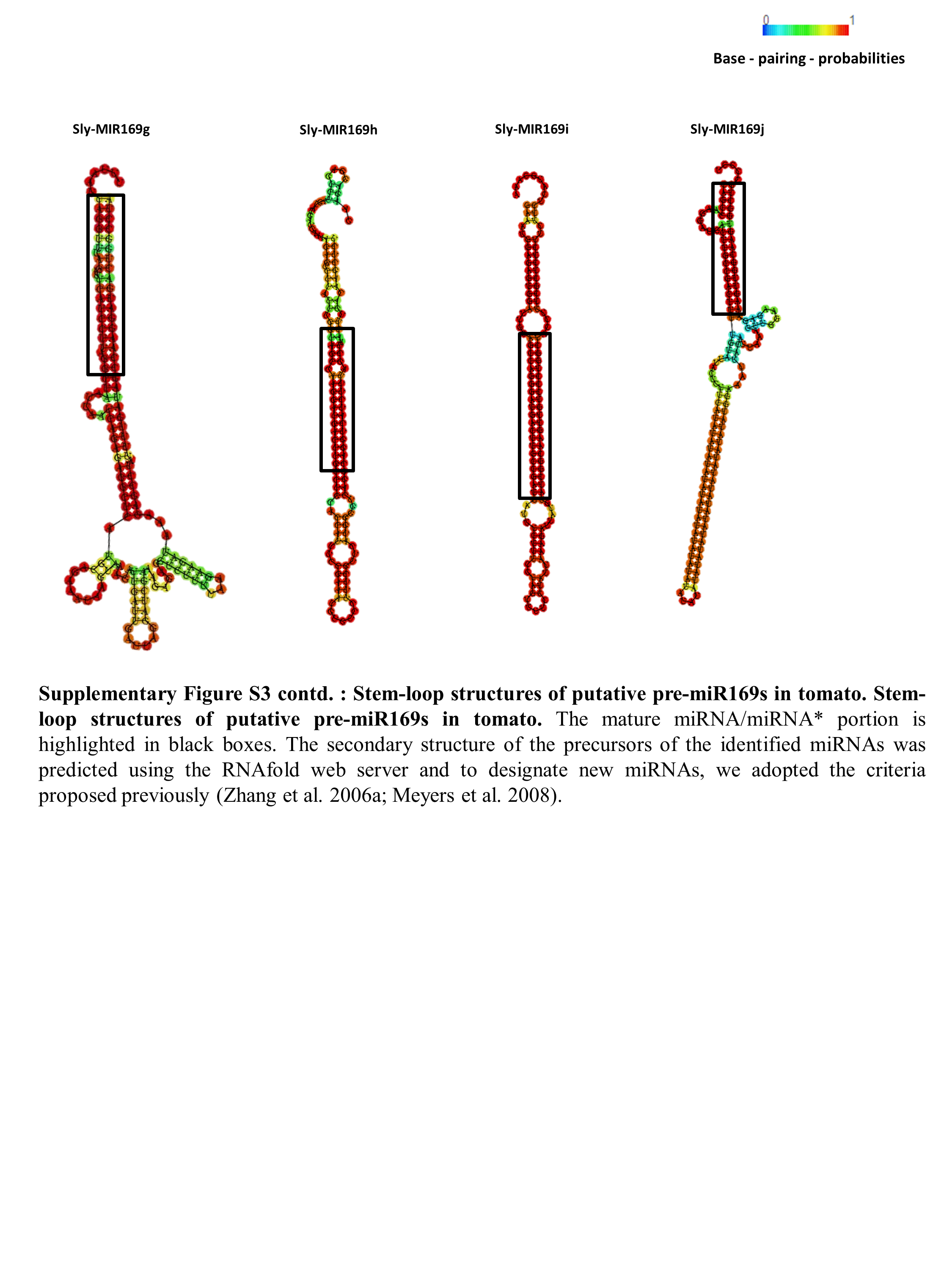

### Supplementary Figure S4: Chromosomal localization, clustering and duplication of MIR169 family genes in tomato.

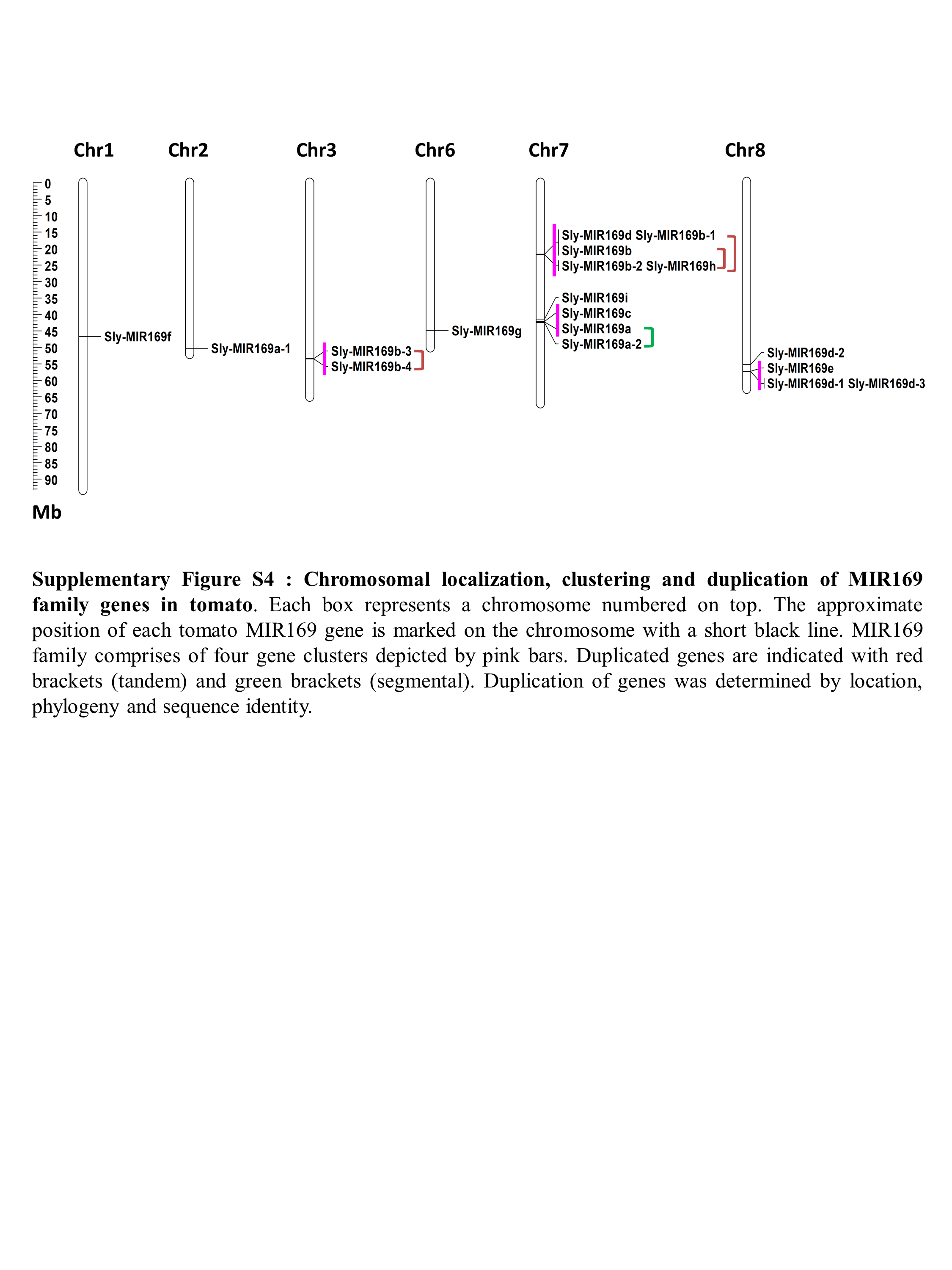

### Supplementary Figure S5: Genomic organization of polycistronic clusters of miRNA169s on chromosome 7

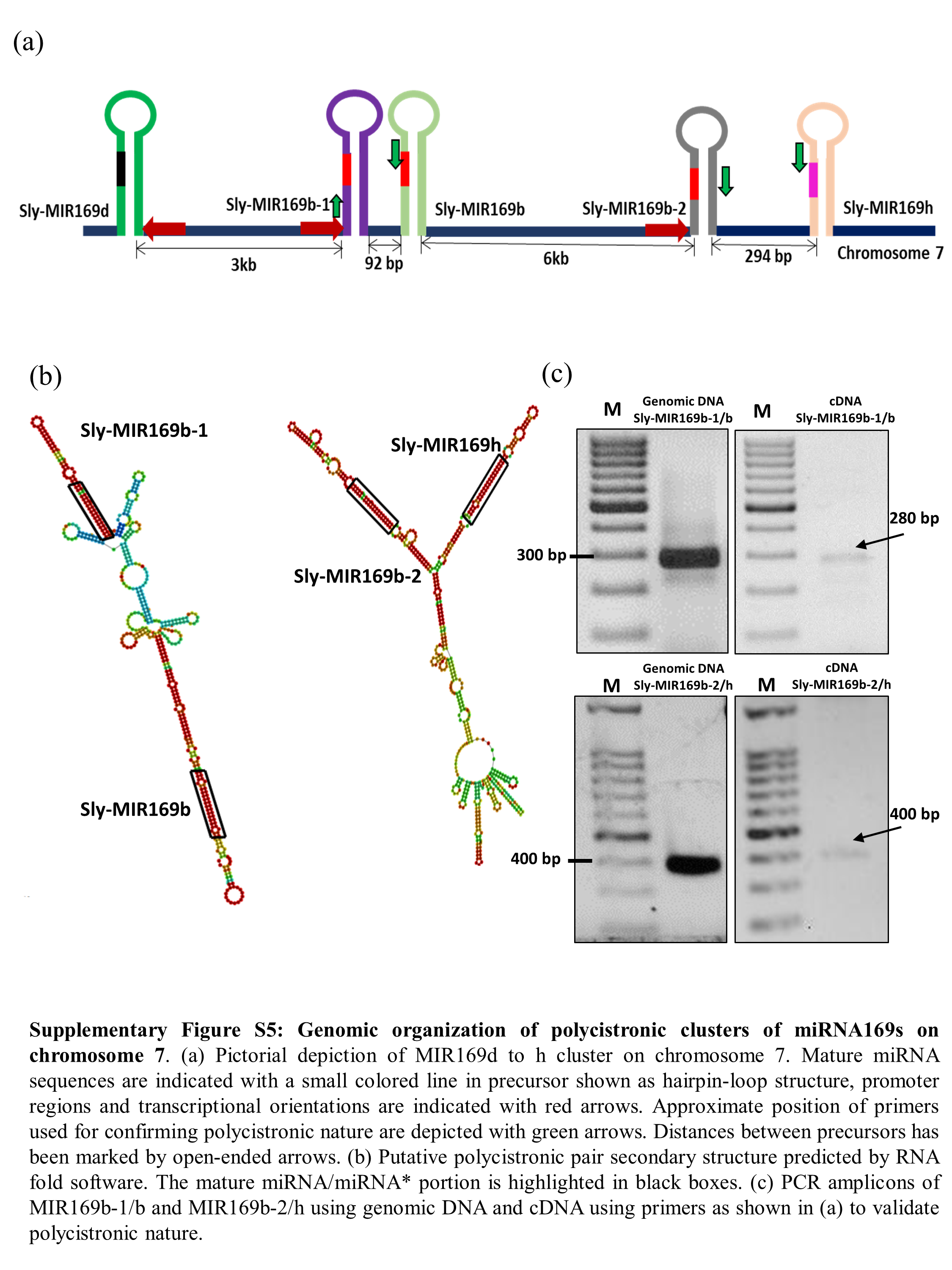

### Supplementary Figure S6: A species-specific MIR169h locus is present in cultivated tomato.

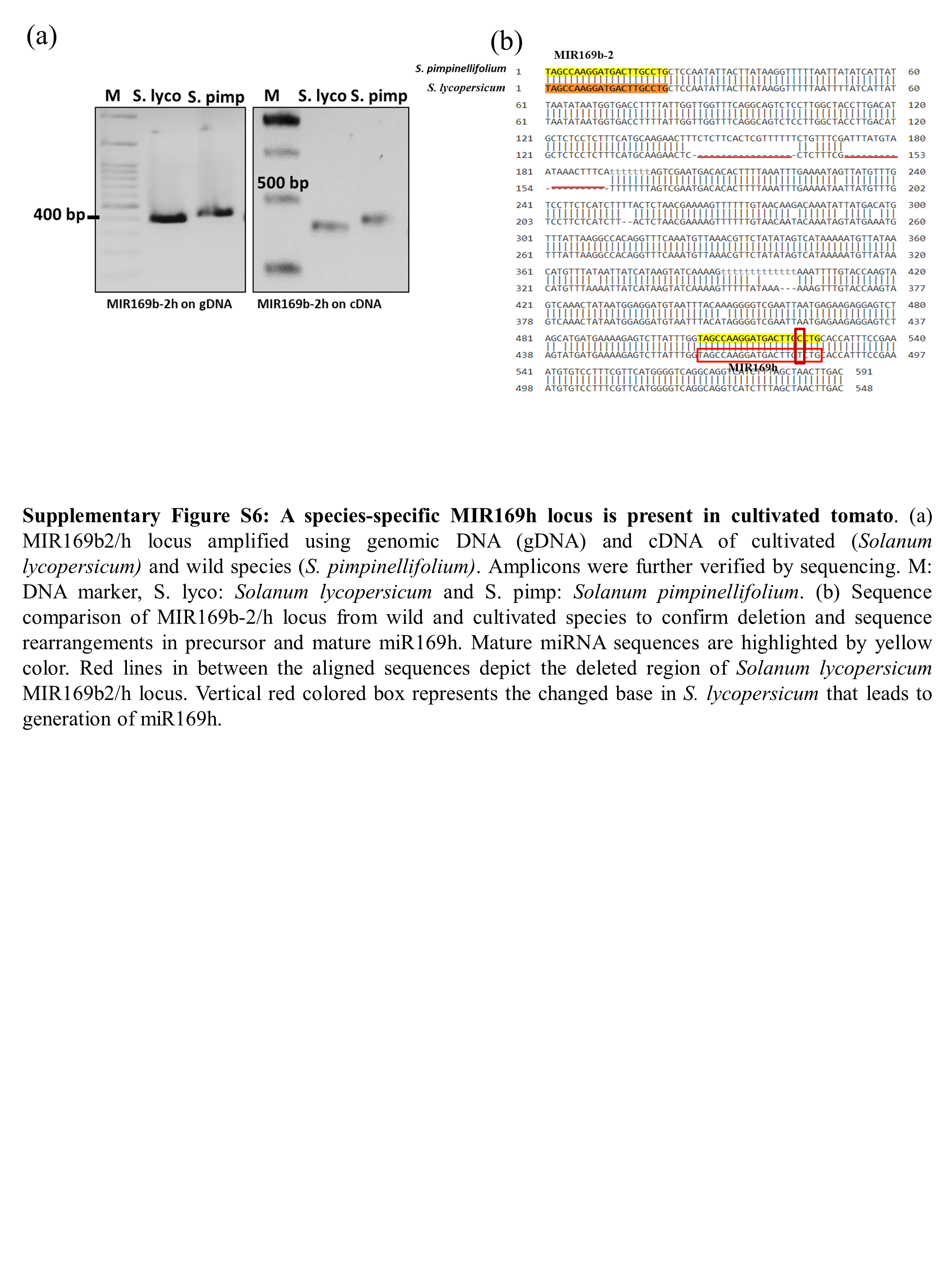

### Supplementary figure S7: Comparison of MIR169b-1/b locus between wild and cultivated tomato species.

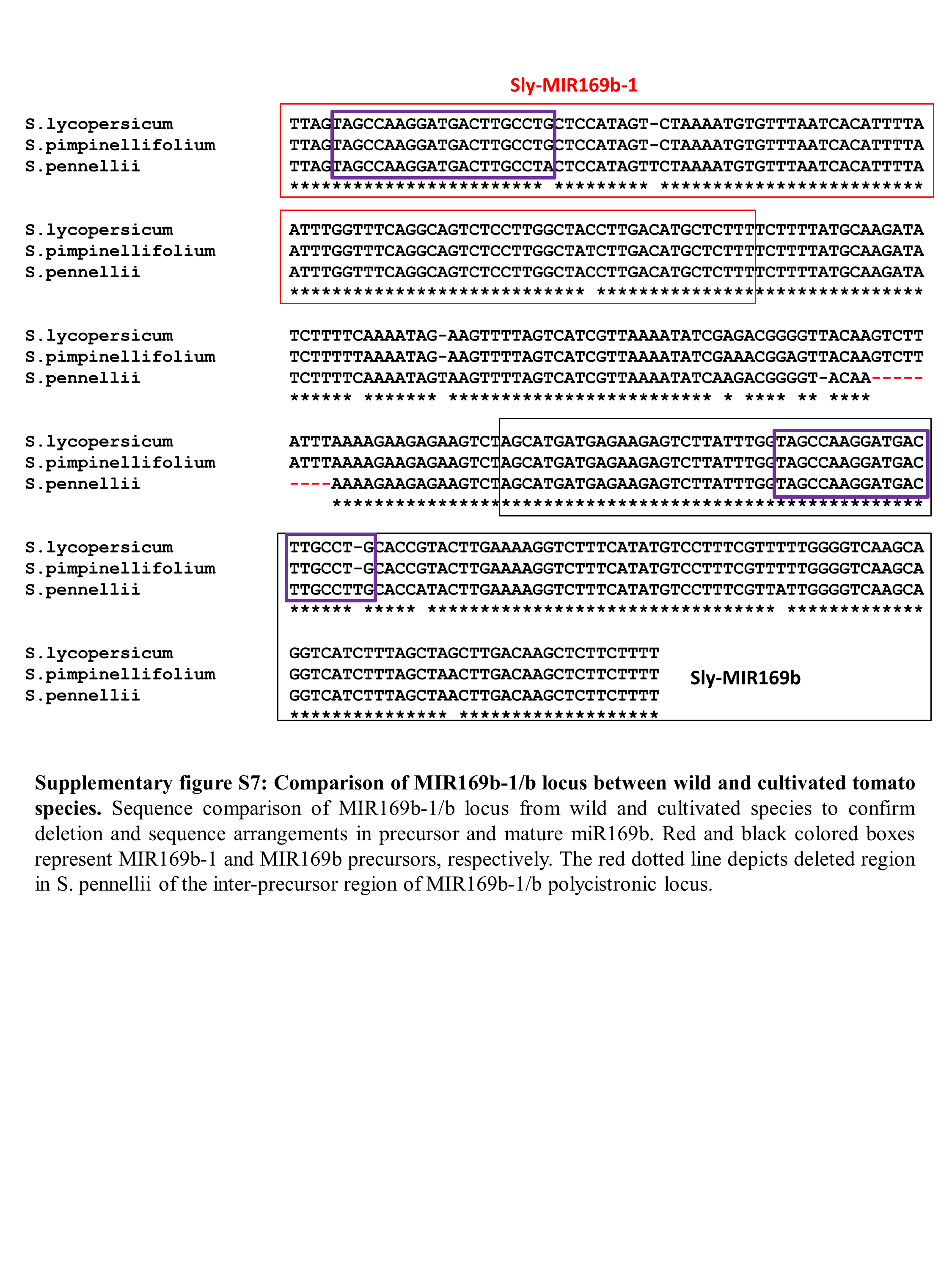

### Supplementary Figure S8: Comparison of MIR169b2/h polycistronic locus between wild and cultivated tomato species.

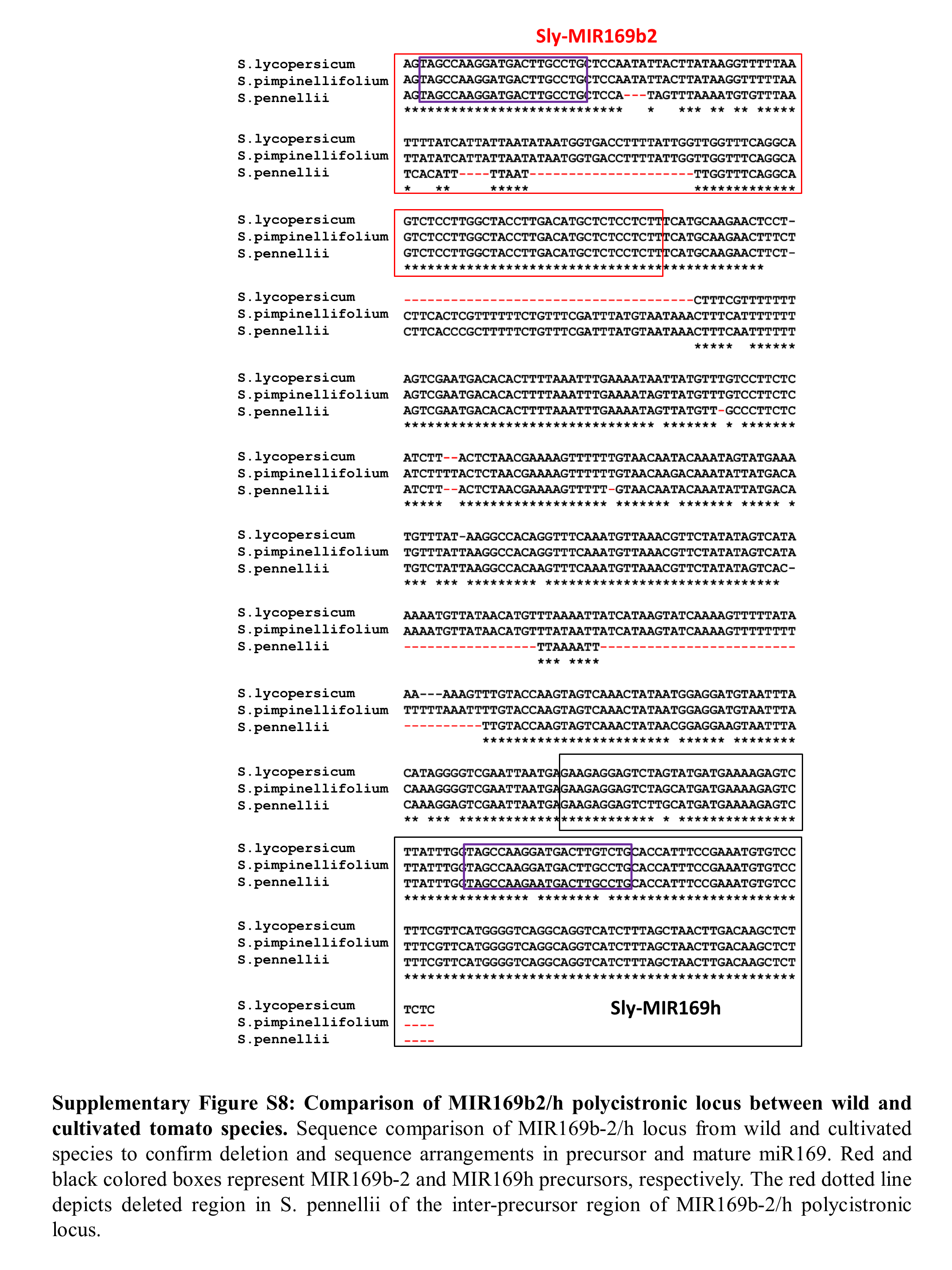

### Supplementary Figure S9: Acquisition of different promoters by MIR169 paralog pairs in tomato.

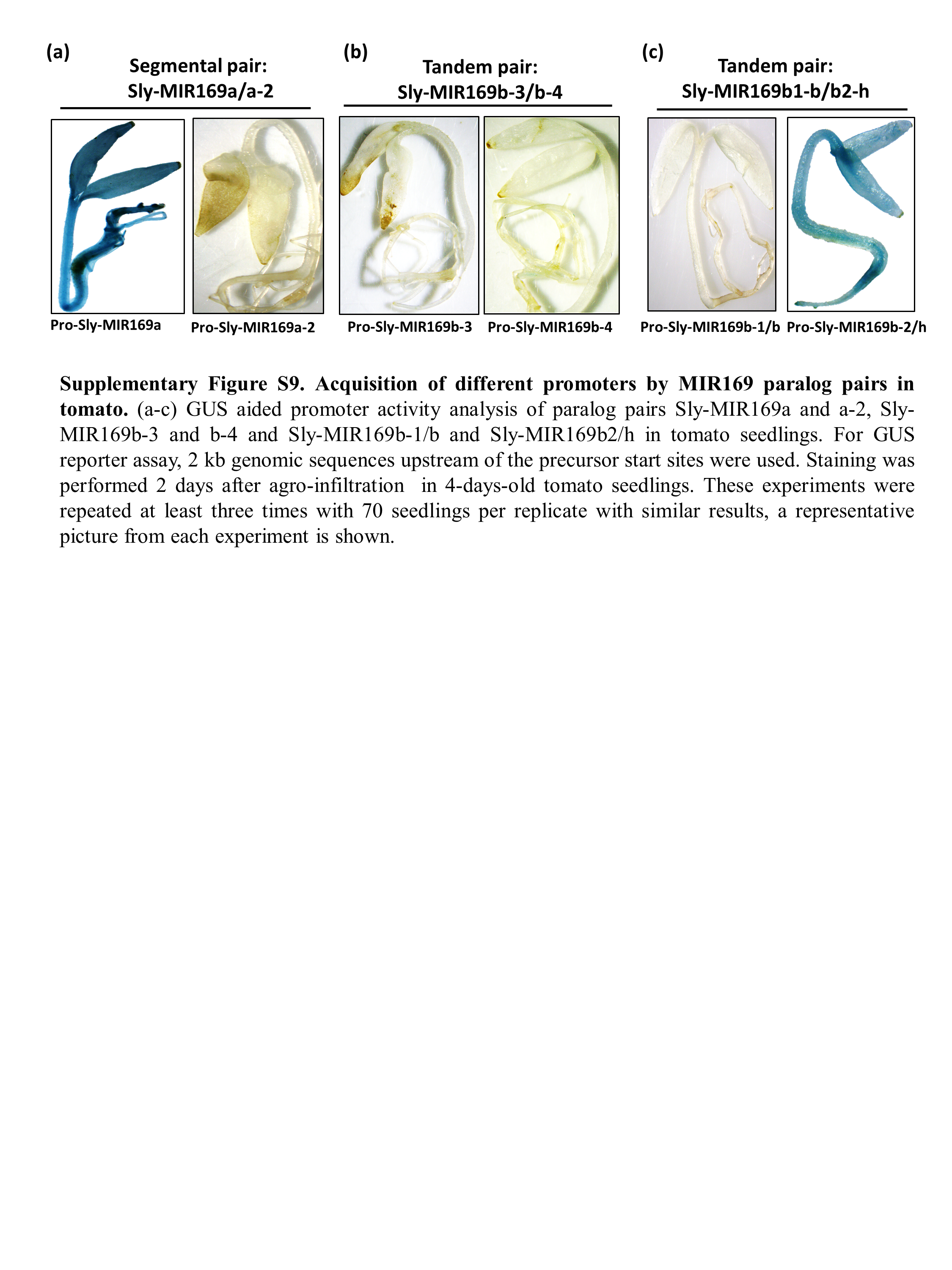

### Supplementary Figure S10: Alignment of novel targets with miR169 in tomato.

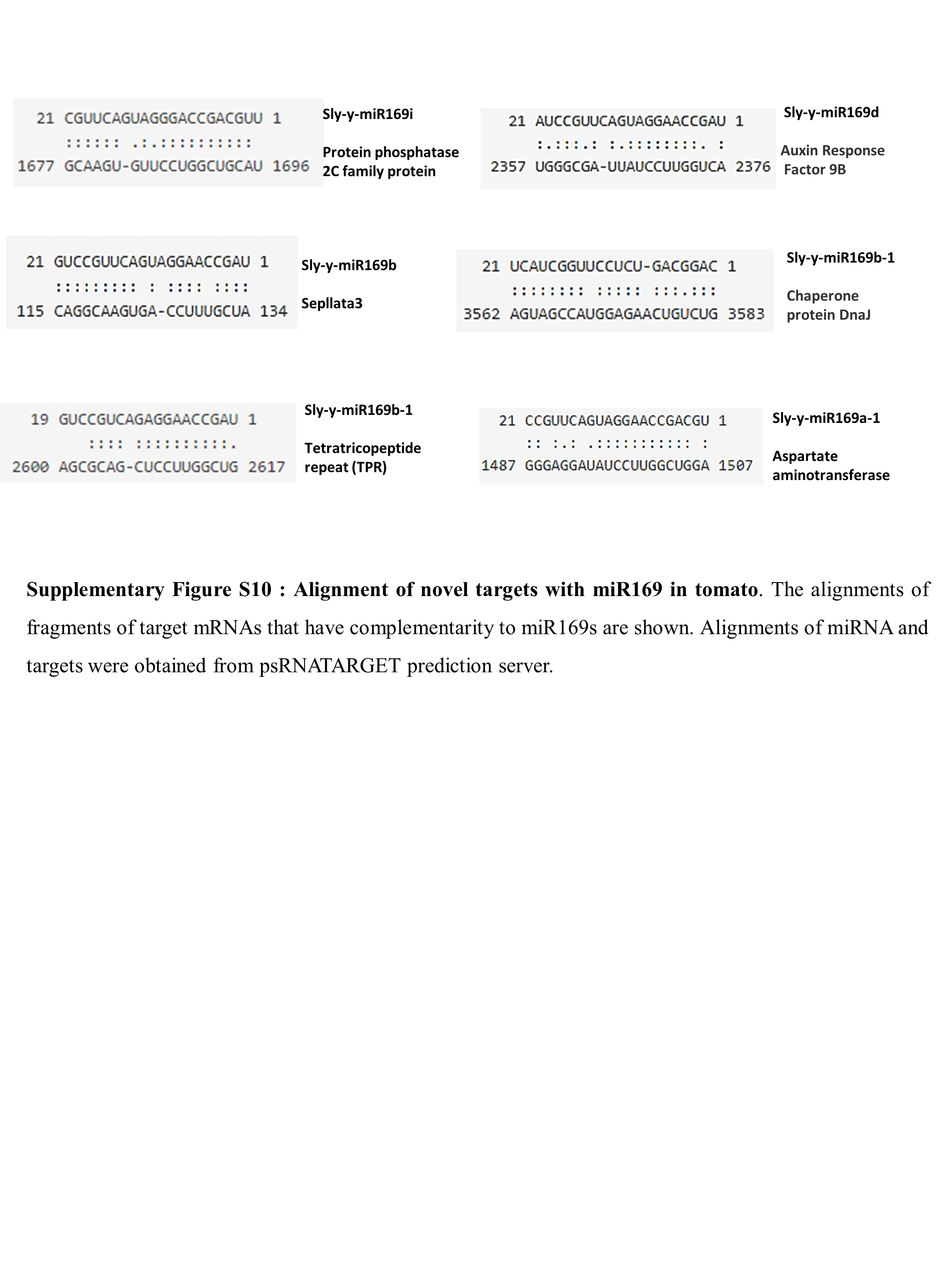

### Supplementary Figure S11: The miRNA169: target functional module on the basis of pathway analysis in tomato.

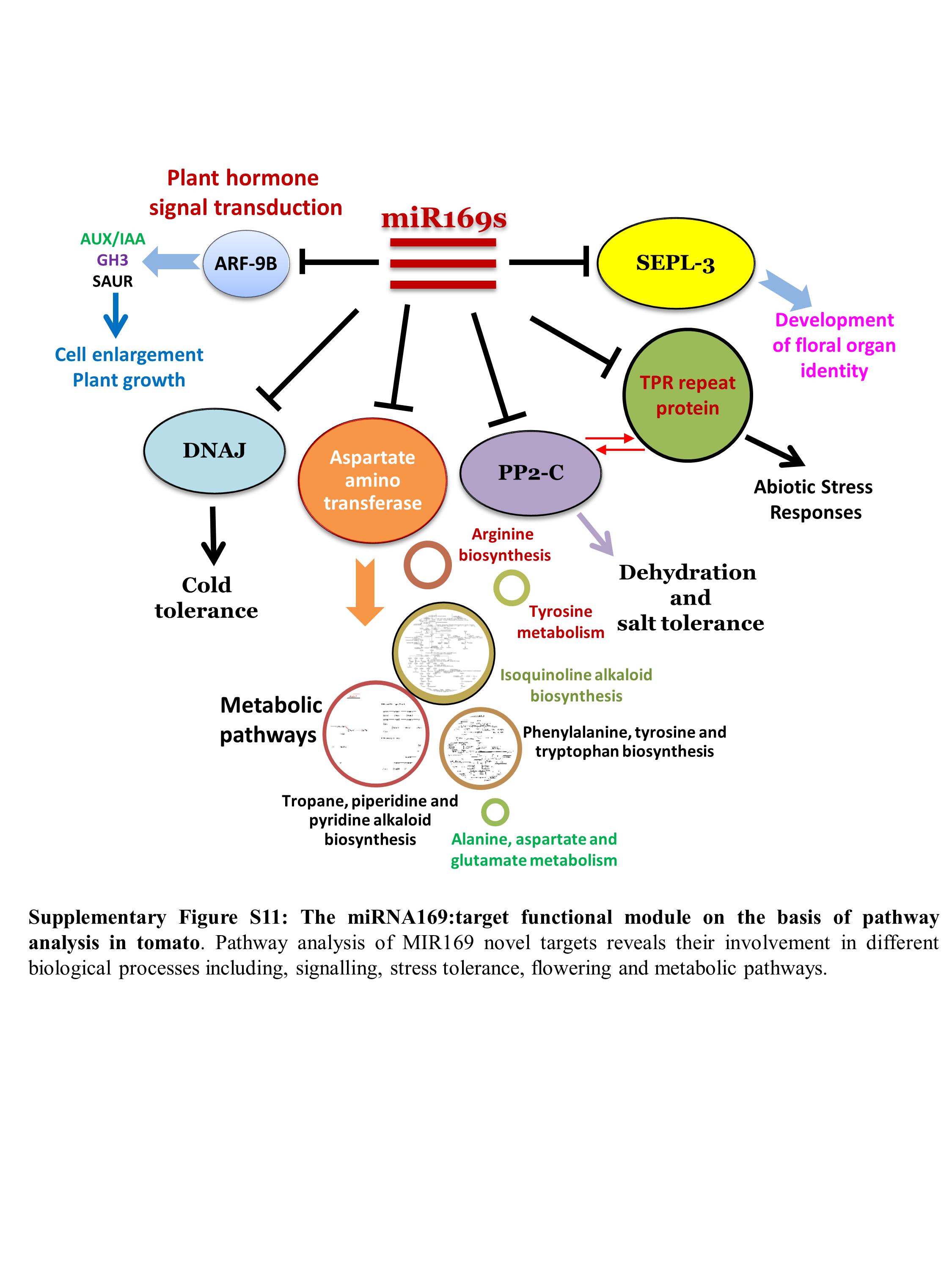
